# Comprehensive profiling of infant gut virome assembly reveals associations with eczema and wheeze

**DOI:** 10.64898/2026.06.24.734081

**Authors:** Sanzhima Garmaeva, Nataliia Kuzub, Asier Fernández-Pato, Sofia Sheveleva, Jody Gelderloos-Arends, Marloes Kruk, Anastasia Gulyaeva, Trishla Sinha, Johanne E. Spreckels, Siobhan Brushett, Cyrus A. Mallon, James A.D. Docherty, Lifelines NEXT cohort study, Edze R. Westra, Jingyuan Fu, Alexander Kurilshikov, Alexandra Zhernakova

## Abstract

Infancy is a critical developmental window during which the gut ecosystem assembles and helps train the immune system, thereby setting trajectories for lifelong health. Bacteria and viruses are equally numerous in this early ecosystem, yet the gut virome’s composition, dynamics, and health relevance remain poorly understood. Here, we show that the infant gut virome is diverse, dynamic, and linked to health outcomes. We performed comprehensive virome profiling of 1,110 longitudinal fecal samples from 314 mother–infant pairs from the Dutch birth cohort Lifelines NEXT using both virus-like particle enrichment (VLP) and total metagenomic sequencing (MGS). We find only 18.9% compositional overlap between the VLP- and MGS-metaviromes, with VLP recovering the active virome and most novel species and MGS predominantly capturing temperate phages. By combining both methods, we identified 8,348 novel virus species spanning diverse hosts, from bacteria to humans, and all major viral genome types (dsDNA, ssDNA, and RNA). We find that bacteriophages frequently encode metabolic functions, including genes related to B vitamin metabolism. We further observe that the development of the infant gut virome is shaped by both host factors, including delivery mode and feeding practices, and continuous switching of temperate phage lifecycles. Notably, the relative abundance of induced temperate phages is also associated with eczema development within the first year of life. Together, these findings establish the infant gut virome as a dynamic and clinically relevant component of early-life microbial development and highlight how comprehensive dual-method profiling is a necessary framework for future virome research.

## Introduction

The early-life gut microbiome is a key determinant of individual health^1,2^. Factors such as mode of delivery, feeding practices, and antibiotic exposure have been shown to strongly influence microbiome composition, with consequences for immune development and later health outcomes^3,4^. While temporal and inter-individual variation in the bacterial gut community has been profiled at large scale^1,5,6^, the gut virome—which constitutes a substantial fraction of the gut microbiome—remains comparatively under-studied. Viruses are at least as numerous as bacteria in the human gut^7,8^ and play a central role in shaping bacterial communities through predation, horizontal gene transfer, and modulation of bacterial fitness^8–10^. Nevertheless, only a limited number of small-scale studies have specifically investigated the gut virome in the first months of life^11,12^. These studies indicate that the early-life gut virome forms a highly individual-specific community^9,12^ that contains predominantly bacteriophages^13^. Infant gut viruses originate at least in part from induced prophages, temperate phages integrated into bacterial genomes^11^ that are transmitted from the maternal gut^14^. Once induced, these prophages produce free virions, with temperate phages dominating the infant gut virome^9,14^. A larger cross-sectional virome study in the Danish COPSAC cohort that included nearly 650 1-year-old infants identified associations between early-life viral signatures and later development of asthma, independent of bacterial effects^15^. This finding underscores the potential importance of the virome as an additional, distinct layer of microbial influence on child health. Despite these initial insights, several key aspects of the infant gut virome remain poorly characterized. These include its longitudinal dynamics at scale and at high temporal resolution, in particular the lysogeny-induction balance of temperate phages and the persistence of early-life colonizers and their functional capacity beyond predation on microbial hosts. Resolving these gaps is essential for clarifying how the early-life virome contributes to child health and disease.

However, characterizing the human gut virome remains methodologically challenging, and most studies rely on one of two sequencing strategies: targeted enrichment of virus-like particles (VLPs) or total metagenomic sequencing (MGS) of the entire microbial community. Extracting VLPs and sequencing their genetic material remains the gold standard for studying the active fraction of the virome, represented by free virions^16^. However, this approach requires handling extremely low biomass samples^17^ and complex purification protocols^18,19^, which often limits the study sample-size. In recent years, the field has benefited from the use of total metagenomes to expand viral catalogs^20,21^. However, because many gut phages are temperate and exist as integrated prophages^22^, total MGS struggles to distinguish active viral replication from integrated phage sequences. Additionally, the MGS approach is mostly limited to dsDNA viruses due to technical limitations in recovering ssDNA and RNA genomes^18^. While the performance in virus recovery from VLP vs MGS has been described in a limited number of adult and infant gut samples^16,23^, it remains unclear to what extent total metagenomics accurately reconstructs the active, encapsulated virome at population scale.

To address these gaps, we performed dedicated VLP isolation and sequencing on 1,110 samples from mothers and their infants across the first year of life in the Dutch Lifelines NEXT cohort^24,1^. In this study, we investigated the developmental dynamics of the early-life gut virome and its influence on infant health. By utilizing paired fecal samples, we assessed the capacity of MGS to reconstruct the active viral fraction compared to VLP; established a framework combining both approaches, termed holovirome; comprehensively profiled the maternal and developing infant gut virome spanning dsDNA, ssDNA, and RNA viruses; and annotated novel gut viruses. Finally, we investigated determinants of early-life virome composition and identified associations between virome features and health-related outcomes, providing new insights into the gut virome’s role in early life.

## Results

### Study design: concurrent virome annotation in VLP and MGS samples

To study the developing early-life virome, we used a selection of 1,110 fecal samples from 314 families (326 infants, including 12 twin pairs and 7 sibling pairs, and their mothers) from the mother–infant cohort Lifelines NEXT^24,1^. Samples from infants were collected at months 1, 3, 6, and 12 after birth. Matched maternal samples were collected at month 3 postpartum or at another timepoint if month 3 was unavailable (Figure 1a, Supplementary Data 1). The study cohort consisted of a representative population of predominantly term-born infants. Infants had a mean gestational age at birth of 39.5 ± 1.7 weeks. 82.2% (n = 268) were born via vaginal delivery, of which 29.5% (n = 79) were delivered at home. The extensive information on feeding, delivery factors, environment, and infant health collected is described in Supplementary Data 2.

**Figure 1.**
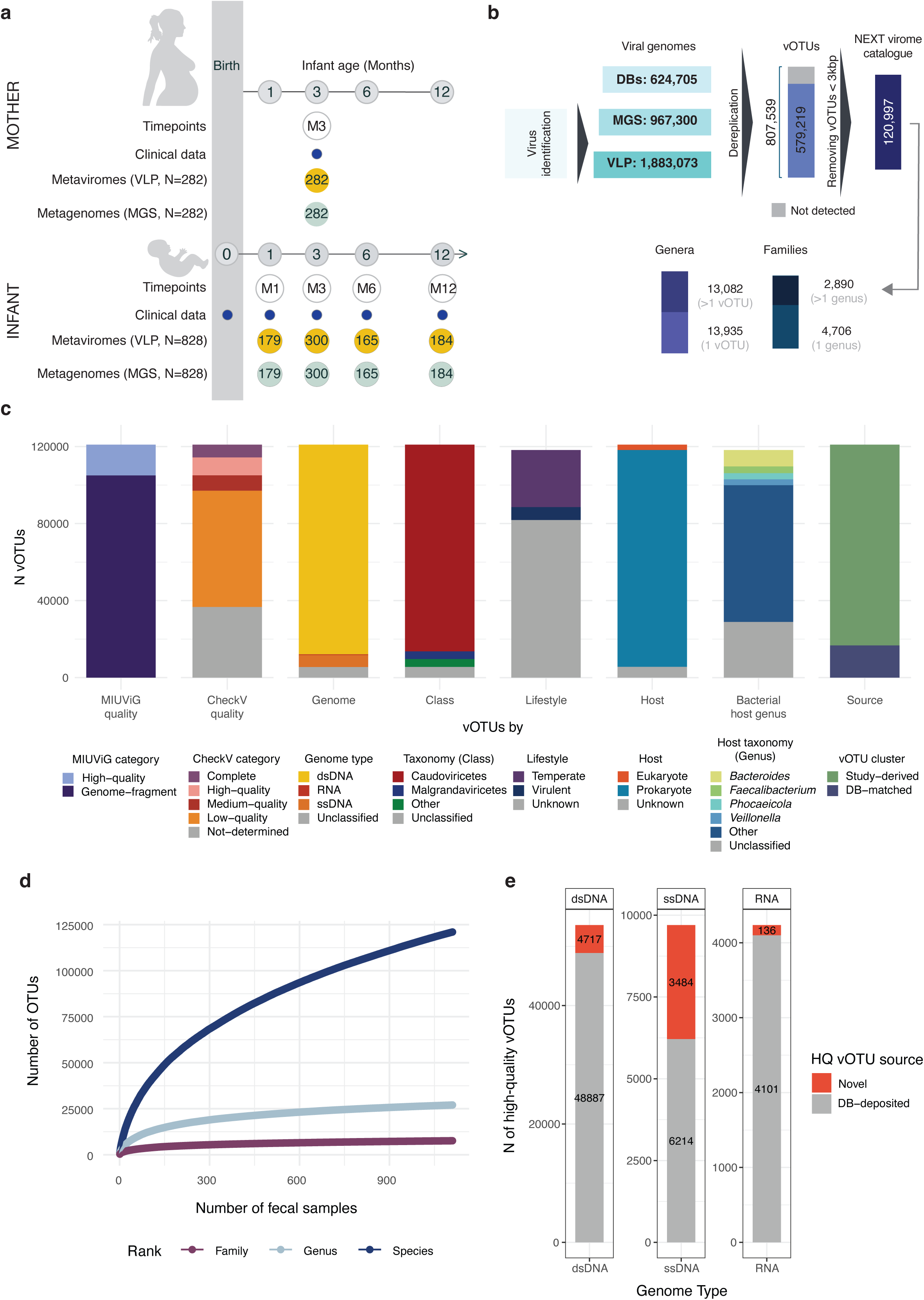
Study design and the NEXT virome catalog. a. Schematic overview of the study design. Fecal samples were collected longitudinally from infants at 1, 3, 6, and 12 months of age (M1, M3, M6, M12) and from their mothers at a single timepoint (M3). For each sample, multiple data layers were generated, including virus-like particle (VLP)-metaviromes, total metagenomic sequencing (MGS)-metaviromes, and matched clinical metadata. The number of samples available at each timepoint is indicated within the colored circles: VLP (yellow) and MGS (teal). b. Simplified overview of the bioinformatic workflow used to build the NEXT virome catalog. Details of read processing, virus identification, and downstream filtering are provided in the Methods. Publicly available viral databases used for dereplication are listed in Supplementary Data 31. c. Characterization of the NEXT virome catalog. The eight bar plots summarize the vOTU composition of the catalog with respect to genome quality (MIUViG and CheckV categories), genome type, taxonomy at class level, predicted lifestyle (bacteriophages only), host domain, bacterial host genus (bacteriophages only), and sequence source (novel: no match in external databases, described: matched to external databases or composed only of external database sequences). d. Accumulation curves for vOTUs, genera, and families across samples. The cumulative number of vOTUs, genera, and families detected (y-axis) is plotted against the number of fecal samples (x-axis), with the VLP- and MGS-detected vOTUs merged per sample. Curves represent the mean across 1,000 random permutations of sample order. Error bars indicate the standard deviation (not visible at the present scale). e. Contribution of novel NEXT vOTUs to existing high quality viral diversity, faceted by genome type (dsDNA, ssDNA, RNA). Each stacked bar shows the total number of high quality vOTUs of that genome type across a combined reference set comprising 11 external viral databases and this study. All high quality vOTUs from the 11 databases were included, irrespective of their detection in NEXT samples.

To comprehensively characterize the gut virome and evaluate the performance of the MGS versus VLP-enriched approaches, we analyzed two complementary datasets derived from the same 1,110 fecal samples: *VLP-metaviromes*, representing the communities of free virions obtained via sequencing of VLP-enriched fecal samples, and *MGS-metaviromes*, representing the communities of all viral genomes present in the samples, including both integrated prophages and free dsDNA virions recovered from total metagenomic data. In addition, we used the MGS data from these samples to study the relations between the virome and the corresponding bacterial community.

### The gut virome is a source for novel viruses

To establish a high-resolution landscape of the human gut virome, we constructed the NEXT virome catalog by integrating sequences across 2,220 paired VLP- and MGS-metaviromes from 1,110 maternal and infant fecal samples. We applied our virome annotation pipeline to this dataset and merged the identified putative viral sequences with viral sequences from 11 external viral databases^21,25–34^ to maximize viral identification (Figure 1b, see Methods). After sequence dereplication and removal of contamination using sequences identified in negative and positive controls and short contigs (< 3 kb), we identified 120,997 viral Operational Taxonomic Units (vOTUs), which were used for downstream analysis (Figure 1b, see Methods). These vOTUs belonged to 27,017 genus-level and 7,596 family-level clusters, indicating that the resulting catalog reflects a wide taxonomic spread of gut viruses and captures substantial diversity.

We next characterized the genomic and taxonomic composition of the NEXT virome catalog (Supplementary Data 3). High quality vOTUs (HQ vOTUs, see Methods) accounted for 13.2% (n = 15,969) of all viruses and ranged from 3 to 396.8 kbp in size (Figure 1c, Supplementary Figure 1a). 88.7% (n = 107,384) of identified viruses were dsDNA *Caudoviricetes* phages, followed by ssDNA *Malgrandaviricetes* phages (3.3%, n = 4,024). The majority of phages with an assigned lifestyle were temperate phages (81.7%, n = 29,625). All phages were predicted to target a phylogenetically diverse range of bacterial hosts, with the highest representation among *Bacteroides* and *Phocaeicola*. Eukaryotic viruses accounted for 2.3% of the NEXT virome catalog, including most of the RNA viruses identified (423 out of 435). Hosts of eukaryotic viruses spanned vertebrates (47.1% of all eukaryotic viruses, n = 1,318), protists (40.5%), and plants (6.7%). Overall, the NEXT virome catalog encompasses broad viral diversity across genome types, taxonomic groups, and lifestyles, providing a representative cross-section of the major constituents of the human gut virome.

Notably, 86.2% (n = 104,247) of the NEXT virome catalog consisted of study-specific sequences not found in external databases (Figure 1c), and 37.5% of these study-specific sequences originated from infant samples (Supplementary Data 4). Together with the observed sub-linear power-law increase in the number of identified vOTUs with increasing sample size (b = 0.46, Figure 1d), this highlights the large inter-individual variability of the virome. This observation is also in line with other studies that found that the majority of viruses identified in infants and adults are individual-specific^9,35^ and that no virus diversity saturation is attainable at the vOTU level^31^.

Of the study-specific vOTUs identified in the NEXT virome catalog, 8,348 (6.9%) were HQ vOTUs that could be categorized as novel viral species, together expanding viral taxonomy by 439 genera and 116 families. The majority of novel vOTUs (56.5%, n = 4,717) comprised dsDNA viruses. However, 3,484 (41.7%) were ssDNA, increasing the diversity of known ssDNA viruses by 56.1% compared with the viral databases we considered^21,25–34^ (Figure 1e). These ssDNA viruses mainly included members of *Malgrandaviricetes* and *Faserviricetes*, two viral classes containing small bacteriophages with circular genomes.

While most of the novel viruses targeted prokaryotes (94.9%, n = 7,919), specifically key early-life bacterial families^1^ such as *Bacteroidaceae* (18.2%, n = 1,517) and *Enterobacteriaceae* (6.1%, n = 510), we also identified 407 novel eukaryotic viruses. These eukaryotic viruses included candidate novel human anelloviruses, animal circoviruses, and human picornaviruses (mainly enteroviruses), with novelty defined operationally by the 95% genome-wide average nucleotide identity threshold used to delineate vOTUs.

All the novel viruses were characterized by a high degree of inter-individual specificity. 59.5% (n = 4,963) were detected in a single fecal sample, and only 11 novel vOTUs (0.13%) were present in >5% of fecal samples. This high proportion of novel, individual-specific viruses once again highlights the human gut virome’s individual-specificity.

### Gut viruses frequently encode diverse auxiliary functions

Given the growing recognition that phages encode auxiliary functions that may influence microbial physiology^36,37^, we investigated the functional potential of viral genomes. Novel and described HQ vOTU representatives (vOTUrs, n = 15,969) encoded 945,727 genes, of which only 35.6% could be functionally annotated (see Methods). Most of these annotated genes (88.9%, n = 299,288) encode structural viral proteins or functions related to transcription regulation, integration, excision, lysis, or nucleic acid metabolism (Figure 2a).

**Figure 2.**
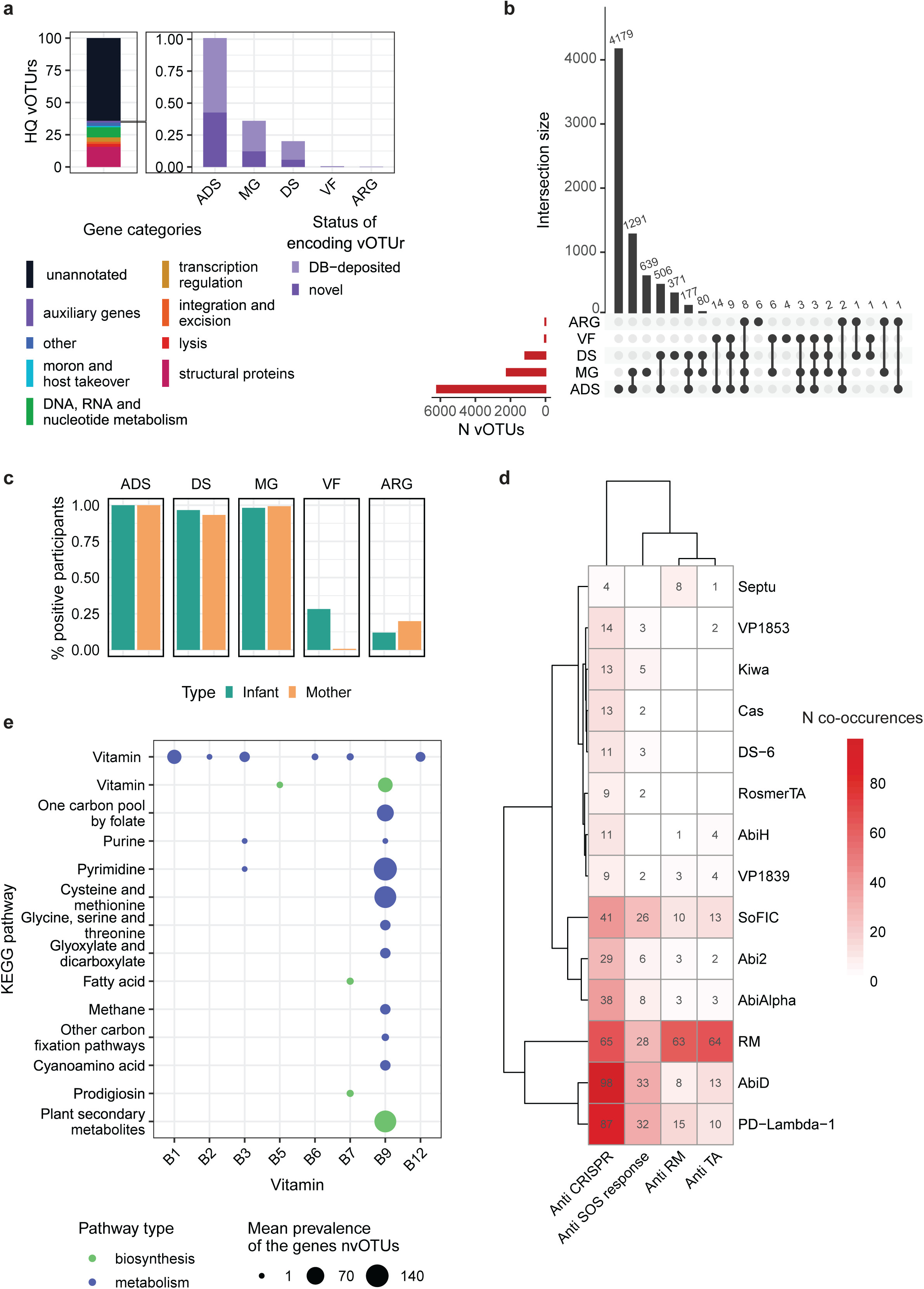
Viral functional capacity beyond the canonical functions. a. Distribution of gene annotations across HQ vOTUrs from the NEXT virome catalog. On the left, functional categories based on PHROG annotations, excluding auxiliary genes identified through separate targeted analyses. On the right, expanded view of the genes belonging to the auxiliary gene category. ADS, anti-defense systems; MG, metabolic genes; DS, defense systems; VF, virulence factors; ARG, antibiotic resistance genes. Gene category names were manually curated after PHROG assignment. b. Distribution of genes with auxiliary functions across the vOTU representatives. The intersection represents the number of viruses that encode multiple auxiliary genes. Abbreviations as in (a). c. Prevalence of vOTUs encoding genes with auxiliary functions, stratified by sample type and color-coded for infants (green, n = 326) and mothers (orange, n = 282). Bar heights indicate the percentage of individuals positive for at least one virus encoding the corresponding auxiliary function. d. Heatmap showing co-occurrence of anti-defense (x-axis) and defense (y-axis) system types within the same vOTUrs. Cell values and color intensity indicate the number of co-occurrences. Rows and columns are hierarchically clustered. e. Balloon plot of B vitamin metabolism genes across vOTUs. Vitamin groups defined by KEGG pathway groups are shown on the x-axis. KEGG pathways are shown on the y-axis. Dots indicate the mean prevalence of genes assigned to each KEGG pathway within the same vitamin metabolism group based on KO identifiers across vOTUs.

To examine auxiliary functions in viral genomes, we performed targeted gene annotation using multiple databases and approaches (see Methods), focusing on viral anti-defense systems, bacterial defense systems, metabolic genes (MGs), antibiotic resistance genes (ARGs), and virulence factors (VFs). Together, these categories accounted for 1.5% (n = 14,524) of all genes and were mainly encoded by temperate dsDNA *Caudoviricetes* (Supplementary Figure 1b).

Anti-defense systems were the most prevalent group of auxiliary functions, with 16 anti-defense system types detected in 38.8% of HQ vOTUrs (Figure 2b, Supplementary Figure 1c) and at least one such vOTU was present in all study participants (Figure 2c). While dsDNA phages prevailed in encoding anti-defense systems, we also identified six anti-CRISPR defense genes in ssDNA phages. We further identified a diverse set of defense systems encoded in HQ vOTUrs (Supplementary Figure 1d), with 4.4% of all vOTUrs encoding both defense and anti-defense systems (Figure 2d).

The second most prevalent group of auxiliary functions was MGs (n = 3,411, Supplementary Figure 1e). Interestingly, 615 vOTUrs encoded genes involved in assimilation of all eight B vitamins essential for the human host (Figure 2e), with genes involved in vitamin B9 biosynthesis most prevalent. While some of the genes involved in vitamin B metabolism also support essential viral functions, we identified other vitamin metabolic genes with no clear role in viral infection. These include CobS and CobT, genes essential for vitamin B12 biosynthesis, which occurred together in 12 viruses (Supplementary Figure 1f), mainly infecting *Enterobacteriaceae*.

Lastly, gut viruses rarely encoded VFs and ARGs (Figure 2b). However, VF-encoding viruses were strongly enriched in infants compared to mothers (OR = 54.9, p-value = 1.9e-25, Fisher’s exact test; Figure 2c, Supplementary Data 5), and most infect *Escherichia* (69.2%, 36 genes; Supplementary Figure 2a). Encoded VFs were diverse and included choloylglycine hydrolase (bile acid hydrolase), previously annotated as a VF that enhances tolerance to bile acid in pathogenic *Listeria monocytogenes*^38,39^. Similarly, we identified 19 temperate viruses encoding ARGs that may confer resistance to different antibiotic classes in their hosts (Supplementary Figure 2b,c). Viruses encoding ARGs were detected in 15.6% of all participants (Figure 2c), with the most widespread a temperate Bacteroides phage carrying the 23S rRNA methyltransferase Erm, a key determinant of macrolide resistance (Supplementary Figure 2d,e).

Overall, more than 40% of the gut phages we considered encoded genes with auxiliary functions that have the potential to shape virus–bacteria interactions, bacterial physiology, and the human host, through diverse mechanisms.

### The human gut holovirome: bridging the gap between VLP- and MGS-metaviromes

To assess the extent to which MGS-metaviromes recapitulate active VLP-detected viromes, we compared the virome profiles derived from 1,110 VLP and MGS paired samples (Supplementary Data 1). The main genomic characteristics, such as the numbers of quality-filtered reads and contigs, were comparable between the two metavirome types (Supplementary Figure 3a, Supplementary Data 6). VLP- and MGS-metaviromes contributed equally to the NEXT virome catalog, with 30.4% of all vOTUs recovered exclusively from the VLP-metavirome and 31.7% recovered exclusively from the MGS-metavirome (Figure 3a). Both methods drastically outperformed the external databases, which contributed only 2.5% of the NEXT virome catalog sequences.

**Figure 3.**
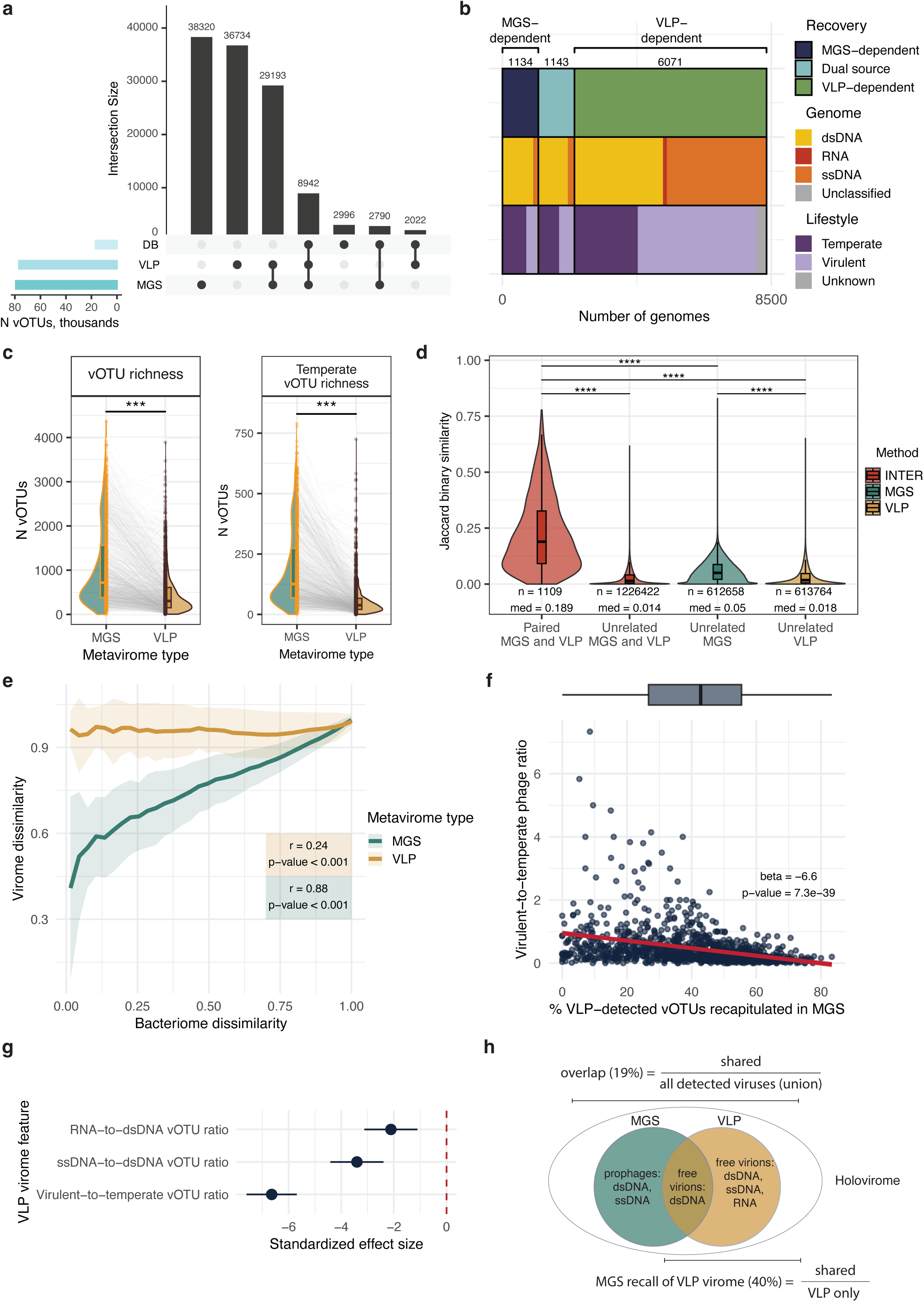
Comparison of MGS- and VLP-metaviromes. a. Contribution of MGS- and VLP-metaviromes and external databases to the NEXT virome catalog (n = 120,997 vOTUs). Intersections show which sources contributed sequences to each post-dereplication vOTU. b. Stratification of novel HQ vOTUs by recovery source. The recovery bar (top) splits vOTUs by the source of the HQ representative of the vOTU. Genome type and Lifestyle bars show the composition of vOTUs within each recovery category. c. Per-sample total and temperate vOTU richness in MGS- and VLP-metaviromes (n = 2,220, 1,110 pairs). Lines connect metaviromes from the same fecal sample. Statistical comparisons were performed using a linear mixed-effect model (LMM). *** p < 0.001. d. Jaccard similarity of vOTU composition between sample pairs colored by comparison type: inter (between MGS and VLP), MGS, and VLP. Statistical comparisons used permutation tests (N = 1,000). *** FDR < 0.001. n, number of pairwise comparisons; med, median Jaccard similarity. e. Concordance between bacteriome and virome between-sample dissimilarity. Bacteriome Bray-Curtis dissimilarities were binned (width = 0.03). The mean and standard deviation of the paired viral Bray-Curtis dissimilarities are shown per bin for both metavirome types. Reported statistics were assessed by Mantel tests. f. Relationship between MGS recall of VLP-detected vOTUs and virulent-to-temperate phage ratio. Red line shows the linear regression fit with 95% CI. The marginal boxplot summarizes the distribution of MGS recall across samples. Statistical significance was assessed using a LMM. All boxplots show the median (center line), 25^th^ and 75^th^ percentiles (hinges), and whiskers extending to 1.5 × the interquartile range. g. Standardized effect sizes (with 95% CI) from LMMs testing whether per-sample VLP virome features explain MGS recall of VLP-detected vOTUs. h. Conceptual definition of the holovirome using a Venn diagram of viruses detectable in MGS, VLP, or both types of the metaviromes (intersection). The outer oval represents the holovirome.

As genome fragmentation can artificially inflate total counts of recovered vOTUs, we evaluated how both methods contributed to the recovery of 8,348 novel HQ vOTUs (see Methods). VLP-metaviromes recovered 3.2-fold more novel HQ vOTUs than MGS, across diverse genome types and lifestyles, with the recovery of 72.7% (n = 6,071) novel HQ vOTUs being VLP-dependent (Figure 3b). While most MGS-exclusive HQ vOTUs (66.5%, n = 754) were temperate dsDNA phages, the majority (53.7%) of the exclusively VLP-recoverable HQ vOTUs (n = 3,261) were ssDNA and RNA viruses that cannot be captured by standard MGS.

To assess how method-specific virus recovery patterns translate to individual samples, we examined the sample-wise composition of vOTUs at the population level. MGS-metaviromes had significantly higher richness than VLP-metaviromes, with a median of 718 (interquartile range (IQR): 384–1,562) versus 305 (IQR: 152–611) vOTUs per sample, respectively (p-value = 5.2e-139; Figure 3c, Supplementary Data 7). While total vOTU richness between the two metavirome types was significantly correlated (beta = 0.25, p-value = 1.01e-16), the difference in richness between paired metaviromes was strongly driven by the differential richness of temperate phages between MGS and VLP (beta = 5.22, p-value < 2.2e-16).

To assess the inter-method consistency, we compared the compositional profiles of MGS- and VLP-metaviromes. MGS- and VLP-metaviromes from the same fecal sample shared much higher similarity than unrelated samples, even when the latter were compared within the same method (FDR < 0.001; Supplementary Data 8, Figure 3d). The overlap between MGS- and VLP-metaviromes accounted for a median of 18.9% (IQR: 9.1–32.6) of all viruses detected per fecal sample using both methods. Both the MGS- and VLP-metaviromes were individual-specific, with very limited overlap between unrelated samples. Notably, VLP-metaviromes showed lower similarity across unrelated samples than MGS-metaviromes (FDR < 0.001), reflecting more divergent and specialized viral recovery in the enriched fraction.

To determine whether the virome provides unique biological information or merely reflects the underlying bacterial community, we compared the strength of co-variation between the bacteriome and the virome structures from VLP- and MGS-metaviromes. We found that the variation in the MGS-derived virome largely followed variation in bacterial community structure (Mantel test, r = 0.88, p-value < 0.001; Figure 3e, Supplementary Data 9), indicating that the MGS-derived virome was nearly a direct reflection of the bacteriome. In contrast, the VLP-derived virome exhibited a much weaker correlation with the bacteriome (Mantel test, r = 0.24, p-value < 0.001).

Given that MGS-detected viromes largely reflected host bacteriome dynamics and had higher temperate phage richness than VLP viromes, we sought to determine the extent to which they captured the active, free-virion fraction. MGS-metaviromes on average captured only 40.4 ± 18.5% of vOTUs identified in the paired VLP-metaviromes (Figure 3f). Recapitulation of VLP-metavirome composition largely depended on the virulent-to-temperate phage ratio, with significantly lower recovery of VLP samples dominated by virulent phages (beta = -6.6, FDR = 7.3e-39; Figure 3f, Supplementary Data 10). In comparison, the number of RNA and ssDNA vOTUs present in the VLP samples had a smaller effect on VLP virome recapitulation (Figure 3g).

Our results demonstrate that VLP- and MGS-metaviromes provide fundamentally different perspectives on the gut virome. While VLP enrichment excels at recovering high quality genomes and broadening the genomic landscape to include novel ssDNA and RNA viruses, MGS remains superior at capturing the integrated dsDNA phage fraction. Crucially, while both methods capture highly individual-specific signatures, they share a remarkably limited per-sample vOTU overlap and represent divergent ecological layers. The MGS-virome strongly correlates with the bacteriome composition, while the VLP-virome is largely independent of it. We therefore propose the concept of the ‘holovirome’ (Figure 3h), which combines data from the VLP-metavirome (the active virome) and the MGS-metavirome (the integrated prophages) to capture the full diversity and dynamics of gut virome.

### The infant gut virome is dynamic and shaped by delivery and feeding modes

We next explored the dynamics of the infant gut virome over the first year of life, focusing primarily on the active virome (VLP), backed up by the holovirome, as the MGS virome has already been described for these samples as part of a larger study^40^. Both the holovirome and active virome richness were lower in infants compared to mothers (FDR < 0.05; Figure 4a, Supplementary Data 11), but both increased with infant age (FDR < 0.05). The infant holovirome expanded faster than the active virome (p-value = 1.4e-04), driven by the number of integrated prophages (p-value < 2.2e-16; Supplementary Figure 3b) and bacterial richness, independent of age (p-value = 4.2e-27; Supplementary Figure 3c). During the first 6 months of life, viral richness was also higher in both the holovirome and active virome of formula-fed infants (FDR < 0.05; Supplementary Data 12, Supplementary Figure 3d,e). These comparisons suggest that the diversification of infant gut viruses is strongly driven by the pace of bacterial community assembly and shaped by infant feeding mode.

**Figure 4.**
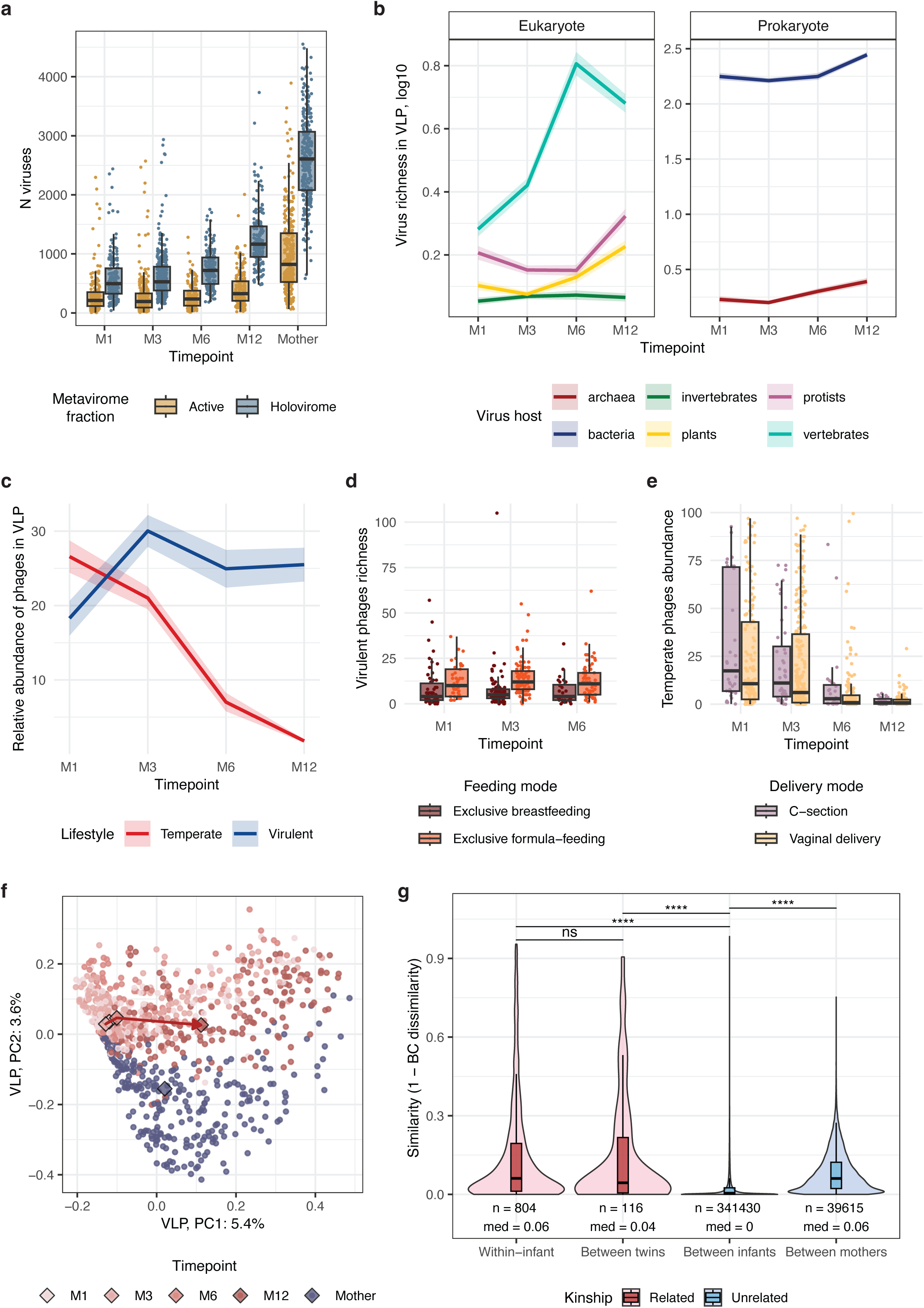
Dynamics and composition of the active virome across early life. a. Active virome and holovirome richness per timepoint in infants (M1, M3, M6, and M12) and mothers (M3) shown as boxplots colored by virome type (active, VLP-only; holovirome, union of MGS and VLP). b. Infant temporal dynamics of vOTU richness in the active virome aggregated by predicted host category. Lines show the mean and ribbons the standard error per timepoint. c. Infant temporal dynamics of the relative abundance of temperate and virulent phages in the active virome. Lines show the mean and ribbons the standard error per timepoint. d. Virulent phage richness in the infant active virome by feeding mode across timepoints M1–M6. e. Relative abundance of temperate phages in the infant active virome by delivery mode across timepoints. f. Principal coordinates analysis (PCoA) of active virome composition based on Bray-Curtis dissimilarity. Each point represents one sample, colored by timepoint. Diamonds mark per-timepoint centroids (medians). Arrows connect infant centroids in chronological order. g. Bray-Curtis similarity (1 − Bray-Curtis dissimilarity) of active virome composition between sample pairs colored by kinship type (Related vs. Unrelated). Statistical comparisons used permutation tests (N = 1,000). *** FDR < 0.001. ns, FDR ≥ 0.05; n, number of pairwise comparisons; med, median Bray-Curtis similarity. All boxplots show the median (center line), 25th and 75th percentiles (hinges), and whiskers extending to 1.5 × the interquartile range.

At the genome-type level, dsDNA viruses dominated both the maternal and infant active viromes (Supplementary Figure 3f). RNA and ssDNA viruses accounted for a small fraction of active virome richness, contributing a median of 0.3% (IQR: 0–0.7) and 2.9% (IQR: 1.5–5.8), respectively, with slight differences between mothers and infants (FDR < 0.05, Supplementary Data 13). Despite this limited contribution to active virome richness, ssDNA viruses reached high cumulative relative abundance in mothers (median 0.32, IQR: 0.13–0.61; Supplementary Figure 3f), revealing a marked decoupling of richness and abundance, consistent with previous reports^35,41^. In infants, ssDNA abundance rose over the first year of life (FDR < 0.05) but remained substantially below maternal levels at every timepoint (FDR < 0.05).

Phages dominated the infant gut virome, while eukaryotic viruses, including viruses of vertebrates and plants, accounted for a median of 1.5% (IQR: 0.6–3.2) of the sample richness. The ratio between the number of eukaryotic viruses and phages was similar between mothers and infants at M1 but diverged from M3 onwards (Tukey p < 0.05; Supplementary Figure 4a, Supplementary Data 14), driven by a significant increase in plant and vertebrate virus richness over time (FDR < 0.05; Figure 4b).

The phage community structure in the active virome was established early in life and remained largely stable across infancy, with similar virulent-to-temperate richness ratios between mothers and infants (Supplementary Data 15). The number of temperate and virulent phages remained stable (Supplementary Figure 4b), while their relative abundances changed throughout the first year of life, in opposite directions (Figure 4c). While virulent phage relative abundance increased with infant age (FDR = 2.5e-03), the relative abundance of temperate phages declined steeply from the peak in the neonatal period (FDR = 2.2e-44).

Feeding and delivery modes also shaped the structure of this phage community. In the active virome, formula-fed infants had a higher number and abundance of virulent phages compared to breastfed infants (FDR < 0.05; Figure 4d, Supplementary Figure 4c, Supplementary Data 12). In the holovirome, the number of temperate phages was also higher in formula-fed infants (FDR = 1.9e-3; Supplementary Figure 4d), reflecting their more diverse microbiome (p-value = 5.9e-5; Supplementary Figure 4e). Delivery mode was associated with the differences in temperate phages: C-section infants’ holoviromes harbored fewer temperate phages compared to vaginally born infants (FDR = 1.0e-3; Supplementary Figure 4f). However, temperate phages were more abundant in the active virome of C-section infants (FDR = 6.8e-34; Figure 4e), suggesting increased prophage induction.

The overall composition of the infant active virome was largely driven by subject identity (R^2^ = 0.51, p-value < 0.001; Figure 4f, Supplementary Data 16), reflecting the high individual-specificity of the early gut virome, as further evidenced by greater compositional similarity within individuals across timepoints and between twins compared to unrelated infant pairs (FDR < 0.05; Figure 4g). Nevertheless, the overall composition of the infant virome changed along a time-defined trajectory (R^2^ = 0.023, p-value < 0.001), becoming more similar among unrelated infants (p-value = 2.0e-47; Supplementary Figure 4g). These longitudinal changes in virome composition were associated with feeding mode and delivery mode (FDR < 0.05; Figure 5a-c, Supplementary Data 17), whereas virome composition was not associated with the presence of pets, maternal smoking history, and family history of allergic disease.

**Figure 5.**
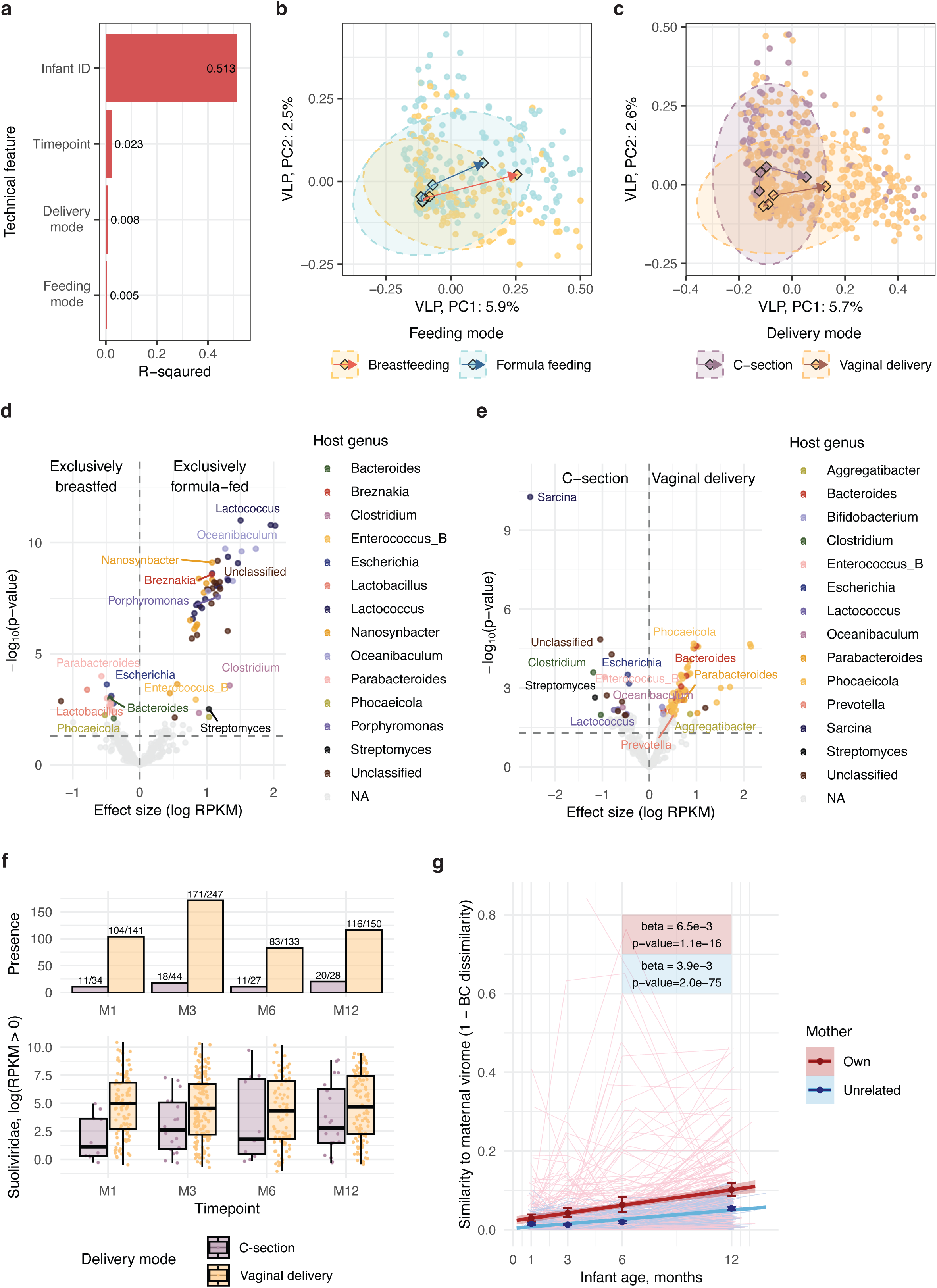
Associations between the infant active virome and early-life exposures. a. PERMANOVA results for host and environmental factors significantly associated with active virome composition. Factors were tested individually with 999 permutations. Bars show the variance explained (R^2^) by each factor. Only significant factors are shown (FDR < 0.05). b, c. PCoA of active virome composition (Bray-Curtis dissimilarity). Each point represents one sample, colored by (b) feeding mode or (c) delivery mode. Ellipses show the 95% confidence region per phenotype. Diamonds mark the per-timepoint, per-phenotype centroids (medians). Arrows connect centroids within each phenotype in chronological order. d, e. Volcano plots of differentially abundant vOTUs in the active virome by (d) feeding mode and (e) delivery mode. Each dot represents one vOTU. X-axis shows the effect size (log RPKM). Y-axis shows the −log10 raw p-value from a LMM. vOTUs significant at FDR < 0.05 are colored by predicted host genus. Non-significant vOTUs are shown in light gray. f. Prevalence and abundance of the *Suoliviridae* family in the infant active virome across timepoints, split by delivery mode. Top, number of infants with *Suoliviridae* detected per timepoint. Bottom, per-sample relative abundance of *Suoliviridae*. g. Bray-Curtis similarity of the infant active virome to maternal viromes over infant age (months), colored by mother type (own vs unrelated). Each infant contributes one similarity value per timepoint per mother type. For the unrelated category, this is the mean similarity to all unrelated mothers. Lines connect timepoints within each infant, showing individual trajectories. Points show the per-timepoint mean across infants. Error bars show the bootstrap 95% confidence interval (1,000 resamples). LMM-derived statistics are reported. All boxplots show the median (center line), 25^th^ and 75^th^ percentiles (hinges), and whiskers extending to 1.5 × the interquartile range.

Specifically, formula-fed infants were enriched for skunaviruses (FDR < 0.05; Figure 5d, Supplementary Data 18), phages that infect *Lactococcus*, likely reflecting phage contamination of dairy-derived formula, as *Lactococcus* itself was rarely detected in infant gut microbiomes regardless of feeding mode (detected in 12 out of 265 samples from formula-fed infants, absent in breastfed infants). Vaginally born infants were enriched for *Bacteroides* and *Phocaeicola* phages (FDR < 0.05; Figure 5e), consistent with the enrichment of *Bacteroides* itself in vaginally born infants (p-value = 4.3e-5; Supplementary Figure 4h, Supplementary Data 19). This pattern was reinforced at the viral family level, with the strongest association observed for phages from *Suoliviridae* (order *Crassvirales*). Phages from *Suoliviridae* were predominantly predicted to infect *Bacteroides* and *Phocaeicola* and were enriched in vaginally born infants throughout the sampling period (log reads per kilobase per million mapped (RPKM) beta = 2.3, FDR = 4.3e-5; Figure 5f, Supplementary Data 20).

Besides feeding and delivery modes, one of the drivers of changes in the infant virome was convergence toward the maternal virome over time (Figure 5g, Supplementary Data 21) and increased vOTU-sharing (p-value = 7.3e-30; Supplementary Figure 5a). Interestingly, infant viromes converged not only to that of their own mothers (beta = 6.2e-3, p-value = 5.9e-45), but also to the centroid of maternal viromes (beta = 3.7e-3, p-value = 2.5e-87), suggesting that infant viromes mature toward a shared adult-like compositional space while remaining highly individual-specific.

Phages infecting *Bacteroides* and *Phocaeicola* were significantly overrepresented among mother-to-infant shared vOTUs (OR = 13.2, p-value < 2.2e-16, Supplementary Data 22) and were the main driver of the increasing number of shared vOTUs (beta = 75.1, FDR < 0.05; Supplementary Figure 5b). This suggests that the maturation of the infant virome toward a maternal-like composition is partly driven by the establishment of *Bacteroides*-infecting phages, consistent with the postnatal expansion of *Bacteroides* in the infant gut (beta = 5.7, p-value = 4.4e-05; Supplementary Figure 5c).

### A personal persistent virome establishes early in life

To further characterize the temporal dynamics of the active gut virome in the early life, we analyzed the VLP-metaviromes from infants for whom we had at least three longitudinal samples (n = 154) using the framework of the personal persistent virome (PPV, vOTUs present in ≥ 3 longitudinal samples) and transiently detected virome (TDV, vOTUs present in < 3 longitudinal samples)^35,41,14^.

We first quantified the total number of vOTUs passing through the infant gut over the first year of life or the cumulative richness over time. Infant gut showed high viral species turnover, with a median cumulative richness of 695 (IQR: 524–1021, Supplementary Data 23, see Methods), which is approximately twice the richness observed at the latest timepoint, M12 (327, IQR: 222–545, FDR < 0.05; Supplementary Figure 5d). Despite this turnover, a small number of vOTUs were detected across at least three timepoints, forming a PPV that contained a median 13 vOTUs per infant (IQR: 5–32). These PPVs accounted for only a median 1.6% (IQR: 0.7%–4.7%) of the cumulative richness over the first year of life per infant (Figure 6a). However, the median cumulative relative abundance of these viruses was 19.2% (IQR: 3.3%–52.7%), although this declined over time (beta = -17.4, p-value = 1.1e-14; Figure 6b). Bacteriophages dominated the PPV composition, with dsDNA and ssDNA phages together accounting for 91.6% (n = 2,683) of all PPV vOTUs. Taxonomic order could not be assigned for the majority of these phages. Of those that could be classified, *Crassvirales* (n = 185) and *Petitvirales* (n = 127) were the most frequent (Figure 6c), in accordance with previous studies^8^.

**Figure 6.**
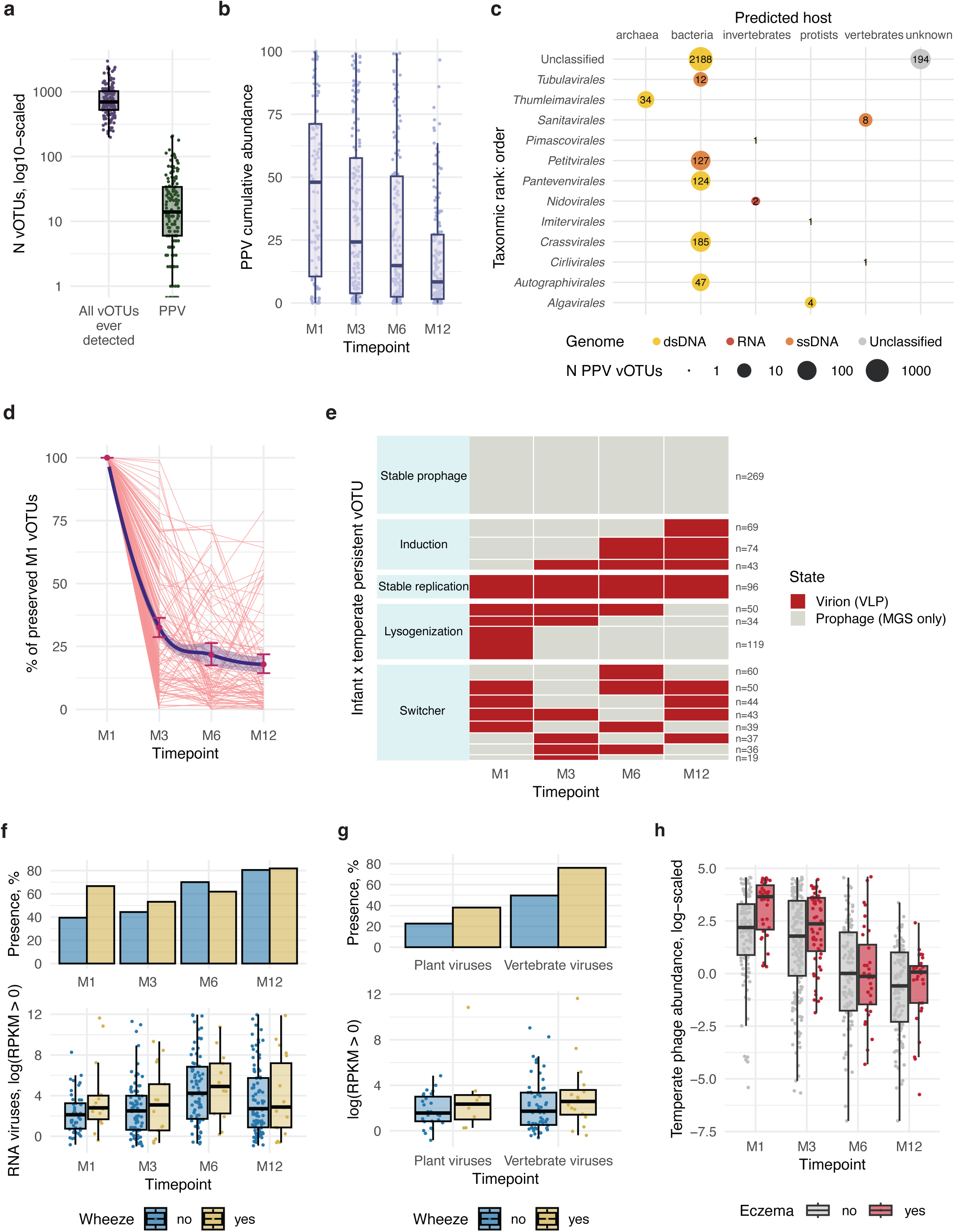
Persistence, turnover, and associations of the infant active virome and health outcomes. a. Per-infant cumulative vOTU richness of the active virome and PPV size. Each dot is one infant. b. Cumulative relative abundance of PPV vOTUs in the active virome per timepoint in infants. Each dot represents a sample, and cumulative abundance is the sum of relative abundances of all PPV vOTUs within a sample. c. Taxonomic and host distribution of PPV vOTUs pooled across infants, shown as a balloon plot. Balloon size and the number inside indicate the count of PPV vOTUs at each host–order intersection. Balloons are colored by genome type. d. Dynamics of M1-detected vOTUs in the active infant virome. Y-axis shows the fraction of M1 vOTUs still detected at each timepoint (M1 = 100% by definition). Lines connect timepoints within each infant. Points show the per-timepoint mean. Error bars show the bootstrap 95% CI (1,000 resamples). e. Prophage–virion state trajectories of temperate persistent vOTUs across infant timepoints. Tile color indicates the detection state at each timepoint: virion (VLP) or prophage (detected in MGS only). Numbers (n) indicate the count of vOTU × infant pairs per trajectory. f. Prevalence and abundance of RNA viruses in the infant active virome across timepoints, split by “ever-wheeze” status (a cross-sectional phenotype reflecting wheeze diagnosis during the 1^st^ year of life). Top, percentage of infants with RNA viruses detected per timepoint. Bottom, per-sample summed relative abundance of RNA viruses. g. Prevalence and abundance of RNA viruses in the infant active virome at M1 split by “ever-wheeze” status. Top, percentage of infants with RNA viruses detected at M1. Bottom, per-sample summed relative abundance of RNA viruses. h. Temperate phage relative abundance in the infant active virome across timepoints, colored by “ever-eczema” status (a cross-sectional phenotype reflecting eczema diagnosis during the 1^st^ year of life). All boxplots show the median (center line), 25^th^ and 75^th^ percentiles (hinges), and whiskers extending to 1.5 × the interquartile range.

We next examined factors distinguishing infant PPV vOTUs from TDV vOTUs. vOTUs detected at M1 were almost 6x more likely to be part of the PPV than the TDV (OR = 5.9, FDR < 2.2e-16; Supplementary Figure 5e, Supplementary Data 24), indicating that persistent viruses enter the gut early. However, this enrichment does not imply a stable early virome. Many vOTUs detected at M1 disappeared rapidly, with only 32.5 ± 25.4% present at M3 (Figure 6d).

Consistent with our previous results for the MGS-metaviromes^40^, maternal sharing and carriage of anti-defense systems increased the likelihood of a vOTU being persistent. Specifically, vOTUs shared with the mother at any timepoint were 5x more likely to belong to the infant PPV (OR = 5.3, FDR < 2.2e-16; Supplementary Figure 5f, Supplementary Data 24). In parallel, PPV vOTUs were modestly enriched for anti-defense and defense systems compared to TDV vOTUs (OR = 1.3 and 1.4 respectively, FDR < 0.05). However, co-encoding of both defense and anti-defense systems, or the presence of other auxiliary functions, did not significantly increase the likelihood of vOTU persistence.

Finally, infant PPVs were modestly enriched in virulent rather than temperate phages (OR: 1.3, FDR = 2.25e-07; Supplementary Figure 5g, Supplementary Data 24), consistent with previous VLP-based study in adults^35^ and our own results from infants’ MGS-metaviromes^40^. However, in MGS-metaviromes of adults, a temperate lifestyle was associated with vOTU persistence^40^, suggesting that a VLP-only PPV definition may undercount temperate phages that undergo lysogenization. To address this, we revisited the temperate vOTUs that were detected in VLP-metaviromes of the given infant (confirming their capacity for induction) but were classified as TDV. For each such vOTU, we then assessed its persistence in the same infant’s holoviromes, capturing the integrated prophage state. Taking these holovirome-persistent lysogens into account increased the PPV size by a median of 9 temperate vOTU per infant (IQR: 3–20), a 69.2% increase over the VLP-only PPV.

We next characterized how frequently temperate phages transition between virion and prophage states during the first year of life. We focused on temperate vOTUs that persisted in the holovirome across all four timepoints within an infant (n = 710 vOTUs, 1,082 vOTU × infant pairs) and classified each such pair by its prophage-versus-virion state across timepoints (Figure 6e, see Methods). A consistent prophage profile across all four timepoints was observed in 269 vOTU × infant pairs, suggesting stable lysogeny. A smaller subset maintained a consistently virion-like profile (n = 96), indicative of active or chronic replication. Interestingly, we also observed dynamic changes between lytic and lysogenic states of temperate phages: in 203 pairs, we observed lysogenization from an initial virion state to a prophage state, and in 186 pairs, the reverse, prophage-to-virion induction was observed. The largest group (n = 328) showed non-monotonic switching between states across timepoints, reflecting repeated cycles of induction and lysogenization. Together, these observations indicate that the highly dynamic infant virome and its viral turnover are shaped not only by the arrival and loss of viral species but also by the lifecycle transitions of temperate phages between prophage and virion states.

### The neonatal virome is associated with health outcomes

We next examined how the infant gut virome relates to health phenotypes, focusing on nine prevalent outcomes, including eczema, wheeze, and cough. For each phenotype, we tested associations between active virome composition and whether the condition was ever diagnosed during the first year (e.g., wheeze recorded only at M12). Details on the phenotype prevalence-based selection, the full list of phenotypes, and diagnostic timing are provided in the Methods and Supplementary Data 2.

The overall composition (beta diversity) of the active infant virome was not associated with any of the tested health outcomes (Supplementary Data 25). We next examined phenotype associations with various features of the active virome, such as viral richness, abundance of temperate phages and eukaryotic viruses (see Supplementary Data 26). We identified several infant virome features at M1 associated with health outcomes. In particular, RNA viruses were more abundant at M1 in infants who developed wheeze later during the first year of life (FDR < 0.05; Figure 6f), specifically plant and vertebrate viruses (Figure 6g, p-values < 0.05, Supplementary Data 27).

We also found that total bacteriophage abundance was lower at M1 in infants who later developed eczema during the first year of life (FDR = 0.008, Supplementary Data 27). In addition, the relative abundance of temperate phages was elevated in these infants at M1 (FDR = 0.004, Supplementary Data 28). To assess whether these differences between infants who developed eczema during the first year of life persisted beyond M1, we fitted longitudinal mixed-effects models across all four timepoints. The total-phage difference was specific to M1 and did not persist longitudinally, whereas the elevation in temperate phage abundance in infants who developed eczema was sustained across the first year (Supplementary Data 29, FDR < 0.05; Figure 6h). This longitudinal temperate phage difference remained robust after adjustment for delivery mode, which we had previously identified as a determinant of temperate phage abundance, and family history of allergic disease, a known atopy risk factor (p-value = 0.02, Supplementary Data 30). Together, these results indicate that an elevated abundance of temperate phages is established by M1, which is before clinical eczema in most infants, and is sustained across the first year of life.

## Discussion

The early-life gut microbial ecosystem plays a critical role in human host development^1,2^. However, while the bacterial component has been extensively studied, the virome remains comparatively underexplored^19^. In this large longitudinal study of 1,110 samples from 314 families in the Lifelines NEXT cohort, we characterized the infant gut virome by combining VLP enrichment with MGS, enabling a comprehensive assessment of both the active and integrated viral fractions. Using this approach, we annotated thousands of novel viral species and genera, characterized their functional potential, described the development of the early-life virome and the lifecycle dynamics of temperate phages, and linked viral characteristics to infant health.

Over the past decade, inter-individual variation in the gut virome has been characterized in increasingly large cohorts^42,40,43^, predominantly using MGS data, driving major expansions in cataloged viral diversity^20,44^. However, it has remained unclear to what extent the MGS-annotated virome represents the active, encapsidated virome. Our results indicate that VLP- and MGS-annotated viromes from the same fecal sample overlap by only 18.9%, on average, in the viruses detected per sample. This low overlap is driven by enrichment of virulent phages and non-dsDNA viruses in the VLP-annotated virome, resulting in the MGS-annotated virome recapitulating only 40% of the active, encapsidated VLP-virome. Further, consistent with the dominance of lysogeny in the healthy human gut^22,8^ and the enrichment of temperate phage recovery observed in MGS-metaviromes, the MGS-annotated virome largely reflects the bacteriome community, while the VLP-virome is mostly independent of it (correlation with bacteriome composition 0.80 vs. 0.24). This observation highlights that the VLP-virome provides an additional omics layer, largely independent from the bacteriome. The characterization of the gut virome therefore requires both VLP and MGS, which together yield the holovirome—the union of all detected viruses, resolved into their active (free virion) and latent (integrated prophage) fractions.

Combination of both VLP and MGS-based approaches in our study enabled recovery of novel viruses across multiple genera and families. Consistent with previous comparisons of virus genome recovery performance between VLP and MGS in human gut^23,45^ and environmental^16^ samples, VLP substantially outperformed MGS in recovering a diverse set of novel viruses across different genome types. The novel ssDNA viruses we identified, which were recovered almost exclusively by VLP, increased the known diversity of ssDNA viruses by more than 50%. Although ssDNA viruses constituted a limited fraction of the gut viruses detected per sample, they were abundant in both infants and adults, making them prominent players of the human gut. In addition to DNA viruses, we identified candidate novel RNA viruses of potential clinical relevance, including enteroviruses. Together, these observations expand the known diversity of the early-life virome and underscore the value of profiling viruses across genome types.

Beyond the expanding genomic diversity of viruses, we also explored their functional diversity and found that gut phages frequently encode genes with auxiliary functions. Among annotated auxiliary gene groups, anti-defense systems were the largest group, despite their diversity likely being under-characterized^46^. Notably, multiple gut phages also encoded genes involved in the synthesis and metabolism of all eight essential B vitamins. These genes likely enhance phage replication, but they may also confer a competitive advantage on the bacterial host because dietary B vitamin availability in the colon is limited by upstream absorption^47^. By altering bacterial vitamin metabolism, phages could also indirectly affect vitamin availability for the human “mega-host”, as some B vitamins can still be absorbed in the colon^47^. These findings support the idea that gut viruses may influence microbiome function and the human host not only through predation, but also through gene transfer.

In adults, a small fraction of an individual’s viruses persists for prolonged periods and dominates the active virome^35,41^. We show that a PPV forms within the first year of life, even though its size is smaller than that of adults^35^. Whether a virus persisted in infants across the first year was linked to whether it carried anti-defense systems, maternal sharing, and its presence at month 1, highlighting the critical role of the early-life window in shaping the gut virome. In parallel, we showed that temperate phages continuously shape the infant active virome through frequent induction from a latent prophage reservoir.

In addition to intrinsic viral features such as phage lifestyle and encoding of auxiliary functions, we found that delivery and feeding modes shape the infant viral community, as seen for the bacteriome^1,3,48^. Beyond the expected associations with phages infecting *Bacteroides* and *Phocaeicola*, we also identified an association between delivery mode and temperate phages that differed between the virome fractions. Temperate phages were enriched in the active virome of CS-born infants, but CS-infants had fewer temperate phages in their holoviromes compared with vaginally born infants, indicating increased prophage induction in this group. Feeding mode was also significantly associated with both overall viral composition and virome richness, as reported previously^11,14,40^. Consistent with diet as a driver of infant virome composition, plant viruses increased from month 6 onward, coincident with the introduction of solid food. Vertebrate viruses were similarly detected more frequently over this period, likely reflecting early-life infections^9^.

Phages, which were traditionally viewed as interacting only with bacteria, have recently been shown to influence human immune responses and to interact directly with human cells^49,50^. Here, we identified a link between early-life virome and health outcomes: a higher abundance of temperate bacteriophages during the neonatal period was associated with an increased risk of later eczema. The immune system likely mediates this association, as phages are recognized by Toll-like receptor 9 (TLR9)^51^, which is involved in the antiviral response. For example, TLR9 and interferon-gamma-signaling-mediated exacerbations of colitis were observed in murine models after oral phage administration^51^, and temperate phages are known to be enriched in Crohn’s disease^52,53^. We therefore hypothesize that the temperate phage–eczema association we observe reflects infant immune stimulation, either directly by viral particles or indirectly through bacterial debris released during prophage induction.

A previous study found that virome-associated asthma risk in children is modulated by the human TLR9 rs187084 variant^15^, supporting a direct interaction between phages and the human immune system through CpG DNA sensing. In our study, we identified an association between increased proportions of eukaryotic RNA viruses of plants and vertebrates and wheezing, a predictor of asthma. Although these viral compartments engage different immune receptors (TLR9 for DNA; TLR3, TLR7/8, and RIG-I-like receptors for viral RNA), both can trigger innate antiviral signaling and shape mucosal immunity^54^. Together, these observations point to the early-life virome as a broader determinant of immune-mediated airway disease, with bacteriophages and eukaryotic RNA viruses likely contributing through parallel but mechanistically distinct routes.

A notable feature of the associations with wheezing and eczema is their temporality. The virome signatures we describe were captured during the neonatal period (month 1), well before the onset of eczema or wheezing in the same children. Therefore, the observed differences do not reflect disease-driven changes in the gut environment, but are instead consistent with the early-life virome acting upstream of immune-mediated disease, rather than being a marker of disease. However, temporal precedence alone does not establish causality, and shared upstream factors influencing both virome assembly and disease risk cannot be excluded in observational data.

Taken together, we have developed a large catalog of early-life viruses and shown that combining MGS and VLP data is necessary to comprehensively characterize infant gut virome development. VLP analysis was particularly powerful for the identification of novel high quality viruses, whereas MGS better captured prophage diversity linked to the developing bacteriome. By analyzing the PPV in infants, we identified evidence for viral persistence, lysogeny, and induction, suggesting dynamic interactions between bacteria and viruses in the infant gut. We also show that viruses carry genes relevant to bacterial metabolism and that viral features are linked to infant health outcomes. These findings position the holovirome as a foundational framework for understanding early-life viral biology and its relevance for host health.

## Methods

### Study cohort

The samples used for this study were obtained from the Lifelines NEXT cohort^24,1^. Lifelines NEXT is a birth cohort designed to study the effects of intrinsic and extrinsic determinants on health and disease. The cohort is part of the Lifelines study, which is a three-generation population-based cohort study of 167,729 individuals living in the Northern Netherlands^56^. For the Lifelines NEXT study, 1,450 pregnant Lifelines participants were included and intensively followed along with their partners and children up to at least 1 year after birth. During the study, biomaterials such as maternal and neonatal (cord) blood, placental tissue, feces, breast milk, nasal swabs, and urine were collected from the mother and child at ten timepoints. Additionally, data on medical, social, lifestyle, and environmental factors were collected via questionnaires at 14 different timepoints and through connected devices. The present study is based on the Lifelines NEXT samples, with no prior selection.

Infant phenotypes and clinical data, including health phenotypes, were obtained and processed as described in Sinha et al., 2024^1^. Health outcome–related phenotypes were selected based on their completeness (phenotype data were available for at least 30% of the infants in the study) and prevalence (for categorical cross-sectional phenotypes, 10% prevalence of every phenotype level within infants was required). The detailed information for the health outcome phenotypes used for the cross-sectional analysis, with the diagnoses derived during the first year of life and summary statistics for all phenotypes, is available in Supplementary Data 2.

### Informed consent

The Ethics Committee of the University Medical Center Groningen approved the Lifelines NEXT study (document number METc2015/600). Participants, their parents, or legal guardians provided written informed consent.

### Sample collection

Fecal samples were collected similarly to the previous Lifelines NEXT–based studies^1,14^. Briefly, samples were provided by mothers who collected their feces during pregnancy (at 12 and 28 weeks), very close to birth, and during the first three months after delivery. Feces from infants were collected from diapers by their parents or legal guardians at the following ages: week 2 and months 1–3, 6, 9, and 12 after delivery. Parents or legal guardians were asked to freeze the stool samples at home at -20°C within 10 minutes of stool production. Frozen samples were then collected and transported to the UMCG in portable freezers and stored in a -80°C freezer until extraction of microbial DNA and viral DNA and RNA. In the current study, 1,110 fecal samples were collected from mothers at 3 months postpartum (when not available, samples from week 12 or 28 of pregnancy or from delivery were used) and from infants at 1, 3, 6, and 12 months after birth. All 1,110 fecal samples were used for both MGS and VLP-enriched sequencing (Figure 1a).

Fecal samples used in this study originated from 321 pregnancies in 314 families (7 with two pregnancies enrolled). Each mother contributed one sample per pregnancy (n = 282 successfully sequenced maternal samples). Each infant (n = 326, including 12 pairs of twins and 7 pairs of siblings) contributed a median of two samples (IQR: 2–3), resulting in 828 successfully sequenced infant samples.

### VLP-metaviromes: genetic material extraction

To study the active virome fraction, metaviromic genetic material was extracted from the fecal samples using a custom protocol^57^ (all reagents and kits catalog numbers are provided in Supplementary Data 32). This protocol was developed by integrating and adapting elements from established methodologies^11,58,59^ to suit the specific objectives and constraints of the present study. For that, approximately 0.5 g of fecal material was resuspended in 10 mL of SM buffer (50 mL 1M Tris-HCl pH 7.5, 20 mL 5M NaCl, 8.5 mL 1M MgSO_4_, 921.5 mL H_2_O) by vortexing at maximum accessory speed for 10 minutes. After homogenization, samples were placed on ice for 5 minutes, then centrifuged at 4°C for 45 minutes at 4,110 rcf to remove large particles and bacteria. The resulting supernatant was filtered twice through a 0.45-µm polyethersulfone membrane filter. The fecal filtrate was adjusted to 15 mL with SM buffer and then concentrated using an Amicon® Ultra-15 Centrifugal Filter Unit (100 kDa MWCO) for 40 minutes at 4,816 rcf. The filtrate concentrate was then adjusted to 15 mL with an SM buffer, and the tubes were inverted 30 times to wash the concentrated VLPs. The diluted filtrate was once again concentrated under the same conditions to approximately 250 µl. Next, 200 µL of the filtrate concentrate was used for nucleic acid extraction after overnight storage at 4°C.

Prior to the nucleic acid extraction, cell-free DNA and RNA were removed using enzymatic degradation, for which 200 µL of the filtrate concentrate was mixed with 24 U of TURBO DNase, 40 U of RNase I, and 24 µL 10X DNase buffer (1 mL 100 mM MgCl_2_, 1 mL 500 mM CaCl_2_, 8 mL H_2_O). This mixture was then incubated at 37°C for 1 h while shaking at 400 rpm. Afterward, the mixture was incubated at 70°C for 10 minutes to stop the enzymatic reaction. The resulting VLP concentrate was then subjected to nucleic acid extraction using the DNeasy Blood & Tissue Kit.

To perform nucleic acid extraction, 180 µL of Buffer ATL and 20 µL proteinase K were added to the VLP concentrate. The mixture was briefly vortexed and incubated at 56°C for 12 minutes while shaking at 400 rpm. Next, 200 µL of Buffer AL was vortex-mixed into the resulting mix and incubated at 56°C for 10 minutes. After that, 200 µL of ethanol (96–100%) was added, the mixture was vortexed, transferred to the DNeasy Mini spin column, and centrifuged at 8,000 rcf for 1 minute. The mixture was washed with Buffers AW1 and AW2 according to the manufacturer’s instructions. Nucleic acids were eluted by adding 25 µL of Buffer AE to the DNeasy Mini spin column, incubated at 37°C, and centrifuged for 1 minute at 8,000 rcf. The procedure was then repeated with an additional 25 µL of Buffer AE to increase the yield. dsDNA concentration was determined using the Qubit 1X dsDNA HS kit.

In addition to fecal samples, two SM-buffer-only (no-template) controls were processed in the same way to serve as extraction negative controls.

### Sequencing of VLP-metaviromes

To enable sequencing of the RNA fraction of the extracted metaviromic genetic material, cDNA synthesis was performed using 33 µL of the metaviromic genetic material and reagents from the SuperScript IV First-Strand Synthesis System (all reagents and kit catalog numbers are provided in Supplementary Data 32). The cDNA synthesis reaction was carried out in a 60 µL reaction volume using random hexamers, following the manufacturer’s instructions.

Briefly, 3 µL of Random Hexamers, 3 µL of dNTP Mix, and 33 µL of the metaviromic genetic material were mixed and incubated at 65°C for 5 minutes, followed by a 1-minute incubation on ice. The mixture containing metaviromic DNA and annealed RNA was then combined with the reverse transcription mix, which included 12 µL 5X SSIV Buffer, 3 µL 100 mM DTT, and 3 µL RiboLock RNase Inhibitor. The combined reaction mixture was incubated at 23°C for 10 minutes, then at 53°C for 10 minutes, before inactivation at 80°C for 10 minutes. The metaviromic DNA and cDNA samples were stored at -20°C prior to DNA library preparation.

DNA shearing, library construction using the xGen ssDNA & Low-Input DNA Library Prep Kit (Integrated DNA Technologies), and sequencing on the NovaSeq 6000 platform (Illumina, San Diego, California) with 2 × 150 bp paired-end chemistry were performed at BaseClear in Leiden, the Netherlands. Sequencing included six negative (no-template) controls and seven positive controls (known-template: genomic DNA of *Staphylococcus aureus* subsp. aureus DSM 20231).

Overall, 6.4 Tb of VLP metaviromic sequencing data were generated for the entire dataset. On average, one VLP-metavirome had 38.1 ± 13.0 million raw reads. Two negative extraction controls produced 40.8 and 36.6 million raw reads, while the negative and positive sequencing controls produced 0.2 ± 0.1 and 16.7 ± 13.9 million raw reads, respectively.

### Metagenomic genetic material extraction and sequencing

MGS metagenomes used in this study were obtained from the European Genome-phenome Archive (EGA, accession EGAD50000001187). Total microbiome DNA extraction and sequencing are described in^1,14^. Briefly, approximately 0.2 g of fecal material was processed using the QIAamp® Fast DNA Stool Mini Kit (QIAGEN), automated on a QIAcube with a final elution volume of 100 µL according to the manufacturer’s instructions, with or without ballcone beads beating, by the Institute of Clinical Molecular Biology (Kiel, Germany). In addition to fecal samples, one no-template control was processed in the same way to serve as metagenome extraction negative controls.

The extracted total microbiome DNA was sheared and library-prepped depending on the DNA concentration using the NEBNext® Ultra™ DNA Library Prep Kit or NEBNext® Ultra II DNA Library Prep Kit (New England Biolabs) for Illumina® and sequenced on the HiSeq 2000 or NovaSeq 6000 platform (Illumina, San Diego, California) with 2 × 150 bp paired-end chemistry at Novogene, China or Novogene, UK. On average, MGS-metaviromes had 28.8 ± 5.0 million reads, while the metagenome extraction negative control produced 8.8 million reads.

### Quality control of VLP- and MGS-metavirome reads

The quality control of VLP-metavirome sequencing reads included adapter and adaptase-introduced tail trimming, read quality trimming and filtering, human reads filtering, read error correction, and deduplication. Sequencing adapters were trimmed by bbduk.sh (BBMap v39.01^60^) with parameters *ktrim = r, k = 23, mink = 11, hdist = 1, tpe, tbo*. The adaptase-introduced tails were trimmed from the reverse reads per the xGen ssDNA & Low-Input DNA Library Prep Kit (Integrated DNA Technologies) manufacturer instructions using bbduk.sh with the parameter *ftl = 10*. To match the length of paired-end reads, the last 10 bases were trimmed from the forward reads with the parameter *ftr2 = 10*. Read quality trimming and filtering and human reads filtering were performed using kneaddata (v0.10.0)^61^ with the following parameters: *LEADING:20 TRAILING:20 SLIDINGWINDOW:4:20 MINLEN:50* and human reference genome (GRCh38p13). Read error correction and deduplication were performed using the settings recommended for adaptase-based library preparation kits^14,62^. Briefly, quality-trimmed, human-data-filtered paired-end reads, as well as unpaired reads, were processed with tadpole.sh (BBMap v39.01) with parameters *mode = correct, ecc = t, prefilter = 2,* and clumpify.sh (BBMap v39.01) with parameters *dedupe = t, subs = 0*. The resulting deduplicated reads were used for all subsequent analyses. On average, a VLP-metavirome retained 29.2 ± 10.8 million quality-controlled reads, the two negative extraction controls retained 19.1 and 16.2 million reads, and the positive sequencing controls retained 9.2 ± 7.9 million reads. Negative sequencing controls retained far fewer reads: one retained none after quality filtering and the others retained a median of 6,228 reads (IQR: 5,830–61,439).

Quality control of MGS reads included adapter trimming, read quality trimming and filtering, and human reads filtering, as described above and in a previous study ^1^. On average, a metagenomic sample retained 27.9 ± 5.1 million quality-controlled reads, and the metagenome extraction negative control retained 7.5 million reads. The quality of the raw and quality-controlled reads was visualized with FastQC (v0.11.9) ^63^ and summarized with multiqc (v1.14)^64^.

VLP-metavirome enrichment for non-bacterial sequences was assessed using ViromeQC (v1.0)^65^ as the ratio of bacterial marker alignment in the paired MGS-metavirome to that in the VLP-metavirome. For MGS metaviromes, the enrichment for non-bacterial sequences was assessed as the ratio of bacterial marker alignment in a sample to the median among MGS-metaviromes. Scores were capped at 100, per the ViromeQC default.

### Assembly of VLP- and MGS-metavirome reads

The quality-controlled and deduplicated metaviromic reads were assembled using settings described earlier^41,62^. In short, paired and singleton reads were assembled using SPAdes (v3.15.3) in single-cell mode with no error correction and k-mer sizes 21, 33, 55, 77, 99, and 127^66,67^. The quality of the assembled contigs was assessed using QUAST (v5.2.0)^68^. On average, 55.8 (IQR: 25.1–99.4) thousand contigs were assembled per VLP-metavirome, of which 5.0 (IQR: 2.1–10.4) thousand were larger than 1 kbp.

The quality-controlled MGS-metavirome reads (paired and singleton) were assembled using SPAdes in metagenomic mode^69^. On average, 67.4 (IQR: 36.7–209.2) thousand contigs were assembled per MGS-metavirome, of which 8.4 (IQR: 4.5–21.9) thousand were larger than 1 kbp.

### Putative virus sequences prediction from metaviromes

Assembled contigs longer than 1k bp were screened to identify putative viral sequences using multiple tools: Cenote-Taker3 (v3.2.1)^70^ in the annotation mode with disabled --prune_prophage; DeepVirFinder (v1.0)^71^, retaining sequences with a score ≥ 0.94^72^; geNomad (v1.7.4)^73^ in end-to-end mode with score calibration enabled (database version 1.9); and VirSorter2 (v2.2.4)^74^ with the flag --include-groups "dsDNAphage,RNA,NCLDV,ssDNA,lavidaviridae". Additionally, VIBRANT (v1.2.1)^75^ (with the -virome flag for VLP-metaviromes) was run on protein-coding genes predicted using prodigal-gv (v2.11.0)^73,76^. Contigs predicted as viral by at least one of these tools were further processed using COBRA (v1.2.2)^77^ to improve assembly contiguity and completeness. The resulting viral contigs were subjected to a two-step prophage pruning. First, geNomad was used to trim prophage regions from putative viral sequences, and sequences without detected prophage regions were retained unchanged. CheckV (v1.0.1, checkv-db-v1.5)^78^ was then applied to all sequences (both trimmed and unchanged) for further prophage pruning. Sequences flagged by geNomad as containing terminal repeats bypassed the CheckV pruning step. CheckV was then run on the resulting (pruned) putative virus contigs to assess virus genome quality and completeness.

Next, putative virus sequences were filtered by excluding: (i) sequences below 1 kbp after prophage pruning, (ii) putative chimeras identified by CheckV (sequences longer than expected genome length), (iii) plasmids identified by geNomad, and (iv) sequences containing an equal or higher number of bacterial genes than viral genes.

### Viral sequence dereplication with external databases

#### Pre-processing of sequences from external databases

Virus sequences from all external databases^21,25,27–34^ except for Viral RefSeq^26^ were processed in the same way as putative virus sequences by pruning prophage sequences, excluding short or chimeric fragments, plasmids, and sequences with an excessive number of bacterial genes. Sequences from Viral RefSeq were used as is. Details on the versions, sources, and numbers of sequences in the external databases are available in Supplementary Data 31.

#### Dereplication of quality-controlled sequences

Quality-controlled viral sequences from 1,110 VLP-metaviromes, 1,110 MGS-metaviromes, and all 11 external databases^21,25–34^ were combined. The combined dataset was then dereplicated according to the Minimum Information about an Uncultivated Virus Genome (MIUViG) species-level thresholds ^44^, defined as 95% average nucleotide identity (ANI) over 85% alignment fraction (relative to the shorter sequence). This resulted in a non-redundant virome catalog comprising 807,539 viral operational taxonomic unit (vOTU) representatives.

### vOTU RPKM table generation and decontamination

#### vOTU abundance estimation

To estimate vOTU abundances, quality-controlled sequencing reads were aligned to vOTU-representative genomes using Bowtie2 (v2.5.4)^79^ with the --very-sensitive preset, retaining only aligned reads in the output (--no-unal). SAM outputs were converted to BAM, sorted, and indexed using Samtools (v1.19.2)^80^. Per-vOTU mapped read counts and coverage were then computed using BEDTools (v2.30.0)^81^. To reduce spurious alignments, read counts with a coverage breadth below 75% of a contig’s length at 1× depth were set to zero^82^, resulting in 579,219 vOTUs detected across all 2,220 metaviromes. Final read counts were converted to RPKM and used for downstream analyses.

#### vOTU RPKM table decontamination and denoising

To decontaminate the vOTU RPKM table, a previously proposed strain-level-based pipeline was applied^17^. Briefly, for vOTUs detected in negative extraction and sequencing controls, as well as positive sequencing controls, consensus sequences were reconstructed using inStrain (v1.9.0)^83^, as follows. Sorted BAM files from samples and control samples were processed with the ‘profile’ module (--database_mode, --min_cov 1), and pairwise genome similarity was estimated using the ‘compare’ module, considering only regions with ≥ 0.5× coverage. Strains were considered shared if their aligned regions showed ≥ 99% population ANI^84^. For vOTUs in biological samples that were shared with controls at the strain level, RPKM values were set to zero. To account for differences in sequencing depth, all strain-sharing cases were considered, including those with < 75% genome coverage. The decontaminated vOTU RPKM table was further filtered by removing vOTUs with a genome size < 3 kbp, reducing noise from short genome fragments but potentially affecting the representation of viral families with small-genome members such as Anelloviridae and Circoviridae^85,86^. The final virome catalog contained 120,997 vOTUs ≥ 3 kbp that were detected in at least one sample.

### vOTU taxonomic profiling, lifestyle and host prediction

#### Combining and expanding vOTU taxonomic assignment

vOTU taxonomic assignment was performed using geNomad^73^ and VITAP (v1.3)^87^, with the database MSL37_RefSeq209_IMGVR. Sequences with a valid Viral RefSeq^26^ match were assigned original Viral RefSeq taxonomy, with a defined set of entries reclassified by VITAP. For the remaining sequences, VITAP was used as the default, with geNomad preferred when VITAP lacked resolution below the class level in *Caudoviricetes*, or when geNomad was considered more reliable for specific groups (*Leviviricetes*, *Picornavirales*, *Aliceevansviridae*). When the two tools conflicted, family-level agreement was used to resolve the conflict. To reflect recent International Committee on Taxonomy of Viruses (ICTV)^88^ changes (Master_Species_List_2024_MSL40.v2), obsolete taxonomy assignments were renamed, and only ICTV-approved ranks were retained.

Next, all vOTUs were clustered into de novo genus- and family-level groups as described in Nayfach et al., 2021^31^. Briefly, protein-coding genes (predicted as described above) from each vOTU representative were aligned against each other using DIAMOND (v2.1.8)^89^ blastp (e-value ≤ 1e-5, ≥ 50% query and subject coverage, up to 10,000 hits per query). Viral genera and families were then defined using the network-based clustering approach of Nayfach et al.^31^, in which genome pairs are scored by AAI and shared-gene fraction, and clusters are extracted with MCL at thresholds calibrated against NCBI RefSeq^90^. The consensus ICTV-curated taxonomy was then extended to previously unannotated vOTUs based on their de novo cluster membership. Starting from the genus AAI cluster, the consensus ICTV-curated taxonomy was propagated to unannotated members based on the most frequent lineage within the cluster (with ties broken by the longest representative). Propagation started at the lowest classified rank in the cluster and moved upward, filling only *Unclassified* ranks. vOTUs whose annotations conflicted with the propagator at any rank retained their original taxonomy. For viral taxons known to frequently having taxonomy misassignments or related to the viruses commonly found in the human gut (*Mimiviridae*^91^, *Poxviridae^92^*), manual curation was performed by inspecting BLAST+ (v2.16.0) nucleotide alignments of vOTUrs assigned to these taxa generated against a local copy of the NCBI nucleotide collection database (nt/nr), downloaded in November 2023. Assignments were evaluated based on percentage of identity (>50%), query coverage (>90%), and alignment length (>200 bp), and consistency of the top hits.

Genome type information (dsDNA, ssDNA, RNA) and host domain (bacteria, archaea, vertebrate, etc.) for all vOTUs with assigned taxonomy were sourced from the ICTV Master_Species_List_2024_MSL40.v2.

#### Phage lifestyle assignment

The lifestyle of vOTU representatives (temperate or virulent) was predicted using BACPHLIP (v0.9.6)^93^. vOTUs with a temperate score ≥ 0.5 were classified as temperate. vOTUs with a virulent score > 0.5 and ≥ 90% genome completeness (CheckV^78^) were classified as virulent. With these cut-offs, lifestyle was assigned to one third of all phages (n = 36,279). RNA viruses and eukaryotic viruses (genome type and host assigned as described above) were excluded from lifestyle assignment. Per-sample relative abundance of temperate and virulent phages was calculated by summing RPKM values across vOTUs within each lifestyle category and dividing by the total vOTU RPKM count.

#### Phage host prediction

Host prediction was performed using iPHoP (v1.3.3)^94^ in two modes: with the default database (Aug_2023_pub_rw) and with a database enriched with previously obtained metagenome-assembled genomes constructed from the Lifelines NEXT cohort study^1^ to improve prediction accuracy. Genus-level predictions were merged and subsequently filtered. For general downstream analyses, only the hit with the highest overall score was retained. In cases where multiple genera shared the same top score, subtool-specific scores were used to resolve the assignment, with the following priority order: BLAST, CRISPR, iPHoP-RF, and RaFAH. If ties persisted after applying these criteria, one genus was randomly selected.

### Novel vOTU definition, prevalence calculation, and building a cumulative curve

#### Study-specific and novel vOTU definition

A vOTU was considered study-specific if no member of its 95% ANI cluster originated from an external database. A vOTU was considered novel if it was study-specific and classified as “high-quality” by CheckV^78^ (per MIUViG standards^44^). Genera and families (defined by AAI- and gene-sharing-based clustering as described above) were considered novel if they contained only study-specific vOTUs and at least one novel vOTU. To estimate the contribution of novel vOTUs to the diversity of viruses in the external databases considered, all sequences from 11 external databases^21,25–34^ classified as “High-quality” by CheckV^78^, regardless of whether they were detected in samples from the present study, were included. Genome type information for external database sequences was derived from geNomad taxonomic assignments and matched ICTV entries^73,88^.

To quantify the prevalence of novel vOTUs in the fecal samples, holoviromes were constructed by combining paired VLP- and MGS-metaviromes into per-sample presence/absence profiles.

#### Source contributions to the NEXT virome catalog

To assess the relative contribution of maternal and infant samples to study-specific vOTUs irrespective of metavirome type, each vOTU was classified by the origin of its cluster members as maternal-only, infant-only, or mixed. The infant contribution was then calculated as: (infant-only + mixed vOTUs) / (total study-specific vOTUs).

Similarly, each vOTU in the entire NEXT virome catalog was classified by the origin of its cluster members across the three sources (VLP-metaviromes, MGS-metaviromes, and all external databases), as visualized by UpSet plot (Figure 3a). Relative contributions of VLP-only, MSG-only, and external database–only were calculated as: (source only) / (size of the entire catalog).

#### Cumulative curve for the catalog size depending on the number of samples

To estimate how the viral catalog size grows with the number of unique fecal samples added, holoviromes profiles generated as described above were used. Cumulative detection curves were generated at the vOTU, genus, and family levels: the order of samples was randomly permuted 1,000 times, and at each step (addition of one sample), the cumulative number of unique units (vOTUs, genera, or families) detected was recorded. The mean and standard deviation of the cumulative curves across permutations were used for Figure 1d.

To characterize the saturation behavior of the cumulative detection curve at each taxonomic level (vOTU, genus, family), three models were fitted to the mean curve: (i) a power-law model (y = a · x^b), fitted by non-linear least squares (stats::nls, v4.4.2^95^) with initial parameter estimates from a log-log linear regression, (ii) a four-parameter logistic model (drc::drm, v3.0-1^96^), and (iii) an asymptotic exponential model (y = Asym + (R0 − Asym) · exp(−exp(lrc) · x)) fitted with stats::nls using the SSasymp self-starting function. Model fits were compared using AIC. The marginal detection rate at the maximum sample count was computed as the first derivative of each fitted model.

### Identification of phage defense and anti-defense systems

To identify phage defense–counterdefense systems carried by phages, we analyzed 15,969 high quality vOTU representatives classified according to MIUViG standards^44^. Protein prediction was first performed for this set of vOTUs using PHANOTATE (v1.6.7)^97^. Defense–counterdefense system predictions were generated using DefenseFinder (v2.0.0)^98–101^, PADLOC (v2.0.0)^102^, and dbAPIS (release from 11/19/2024)^103^ together with DIAMOND (v 2.1.8)^89^ and HMMER (v3.4)^104^ (Supplementary Data 33). The defense systems DMS_other and VSPR were excluded from subsequent analyses^105^. For dbAPIS predictions, hits with a bit score < 50, an e-value > 1e-10, or % sequence identity < 50 were discarded, and only the best-scoring hit was retained. Since dbAPIS predicts individual anti-defense genes rather than entire systems, multiple dbAPIS-predicted genes of the same anti-defense type that occurred consecutively within a viral genome were collapsed and treated as a single anti-defense system. Finally, outputs from all tools were standardized using custom scripts and manual curation, and redundant predictions were dereplicated. Anti-defense systems predicted by dbAPIS as “restriction–modification (RM); bacteriophage exclusion (BREX)” were annotated as “anti-RM”. Anti-defense systems predicted by dbAPIS as “CBASS, Pycsar, and CRISPR–Cas (type III)” were annotated as “anti-defense broad”.

For defense systems, additional boundary checks were performed to confirm that the locus was flanked by viral genes, ensuring its localization within the viral region and excluding potential contamination resulting from imprecise prophage pruning or assembly errors^37^. To achieve this, the predicted proteins were further annotated with the PHROG (v4)^106^ database using HMMER (v3.4)^104^ and with KofamScan (v1.3.0)^107^. Viral genes were defined as: (1) genes predicted within one of the following PHROG categories: ‘head and packaging’, ‘tail’, ‘lysis’, ‘connector’, or ‘integration and excision’, (2) genes whose PHROG or KEGG database annotations contained one of the following keywords: ‘portal’, ‘terminase’, ‘spike’, ‘capsid’, ‘sheath’, ‘tail’, ‘virion’, ‘holin’, ‘base plate’, ‘baseplate’, ‘lysozyme’, ‘head’, ‘structural’, ‘phage’, or ‘vir’^75^, which were subsequently manually checked, or (3) genes belonging to an anti-defense system. Only defense systems with confirmed viral flanks were retained for downstream analysis.

### Identification of metabolic, antimicrobial resistance, and virulence factor genes carried by phages

The proteins (predicted as described above) in 15,969 high quality vOTUrs were annotated using KofamScan (v1.3.0)^107^ (Supplementary Data 34). Hits with bit scores below the profile-specific recommended threshold (or lacking such a threshold) were discarded, and only the best-scoring hit per protein was retained. Genes were classified as metabolic if they corresponded to genes in the “Metabolism” category as defined in KEGG (https://www.kegg.jp/kegg/pathway.html). To further explore the functional potential of the viruses identified in this study, the same set of vOTUs was used to predict antimicrobial resistance and virulence genes. Predicted proteins were annotated using AMRFinderPlus (v4.2.5)^108^ with default parameters and VFDB (downloaded on 16/01/2026)^109^, together with DIAMOND (v 2.1.8)^89^. For VFDB-annotated proteins, only hits with 100% coverage, at least 50% sequence identity, and an e-value < 1e-5 were retained. Hits that were simultaneously annotated as “autolysin” using VFDB and “endolysin” using PHROG were discarded from the virulence factor list. Only the best hit was selected for the final annotation. Similar to defense system identification, additional boundary checks were performed for each predicted metabolic, antimicrobial resistance, or virulence gene to ensure its affinity to the viral genome ^37^.

### Bacteriome community profiling from MGS samples

Bacterial community composition was profiled using MetaPhlAn4^110^, as described previously^1^. The MetaPhlAn database of marker genes mpa_vJan21 and the ChocoPhlAn species-level genomic bin (SGB) database (202103) were used^110^.

### Comparison of MGS- and VLP-metaviromes

The numbers of raw, human, and quality-controlled sequencing reads, the numbers of assembled contigs and contigs ≥ 1 kbp, and the enrichment for non-bacterial sequences (defined by ViromeQC as described above) were compared between MGS- and VLP-metaviromes using a separate linear mixed-effects model for each feature. Metavirome type was included as the main predictor using the lmer function (lme4, v1.1-35.5)^111^ with sampling timepoint as an additional fixed effect and individual as a random intercept to account for repeated sampling. Fixed-effect p-values were obtained using Satterthwaite’s approximation (lmerTest, v3.1-3)^112^. Effect size (Cohen’s d) was determined using effsize::cohen.d (v0.8.1)^113^.

#### High quality vOTU recovery in MGS- and VLP-metaviromes

To compare the performance of MGS- and VLP-metaviromes in recovering high quality vOTUs, we considered vOTU clusters with high quality vOTU representatives according to MIUViG classification assigned by CheckV^44,78^. Within each vOTU cluster, we selected the longest sequence contributed by MGS-metaviromes and the longest sequence contributed by VLP-metaviromes as the method-specific candidate representatives, and recorded the MIUViG quality of each. Clusters were then labeled "VLP-dependent", "MGS-dependent", or "Dual source" according to whether the high quality candidate originated from VLP-only, MGS only, or both, respectively.

#### Viral community similarity between MGS- and VLP-metaviromes

Jaccard similarities were computed on vOTU presence/absence profiles for all pairs of samples using function vegdist from R package vegan (v2.6-8)^114^. One MGS-metavirome that contained no detected vOTUs was excluded, together with its paired VLP sample. We used Wilcoxon rank-sum tests to assess whether Jaccard similarities differed between the following sample pair categories: paired MGS vs VLP from the same fecal samples, unrelated MGS vs VLP, unrelated MGS vs MGS, and unrelated VLP vs VLP. Because the number of unrelated sample pairs is combinatorially inflated, statistical significance was assessed by permutation (n = 1000). For each comparison group (contrast), group labels were shuffled and the Wilcoxon test recomputed, yielding a permutation-based p-value. Permutations were performed within individuals to account for repeated sampling. The contrast comparing unrelated MGS–MGS to unrelated VLP–VLP pairs, where pairs are unrelated by construction, was permuted without this restriction. Final p-values were adjusted across all contrasts using the Benjamini-Hochberg method.

#### Bacteriome–virome co-variation in MGS- and VLP-metaviromes

To reflect each method’s standalone output, community profile tables from MGS- and VLP-metaviromes were filtered to remove cross-method-dependent vOTU detections. From the MGS community profile, we excluded vOTUs exclusively recovered from VLP-metaviromes. From the VLP community profile, we symmetrically excluded vOTUs exclusively recovered from MGS-metaviromes. vOTUs with member contributions from both methods or from external databases were retained in both tables.

To assess bacteriome–virome co-variation across the two metavirome methods, vOTU relative abundances were renormalized within the filtered MGS- and VLP-metaviromes and bacterial community composition was represented by species relative abundances. For each data type, sample-pairwise Bray-Curtis dissimilarities were computed (vegan::vegdist)^114^. The strength of bacteriome–virome co-variation was then assessed separately for MGS and VLP using Mantel tests with Spearman rank correlation and 999 permutations (vegan::mantel)^114^. Analyses were restricted to fecal samples with paired bacteriome and virome data (n = 1,110 for bacteriome–VLP virome analysis, n = 1,109 for bacteriome–MGS virome analysis).

#### MGS recall of VLP-detected vOTUs

Recall analysis was performed on the filtered MGS- and VLP-metaviromes described above. The number of vOTUs detected per sample was calculated within the filtered metaviromes, both overall and across genome types (ssDNA, RNA, dsDNA) and lifestyles (temperate, virulent). For each MGS–VLP sample pair from the same fecal sample, we counted the shared vOTUs. Recall was defined as the percentage of VLP-detected vOTUs also found in the paired MGS sample: shared vOTUs / (number of VLP-detected vOTUs × 100).

To test whether VLP-virome composition is associated with MGS recall, three compositional ratios were computed for each VLP sample: the virulent-to-temperate phage ratio, the ssDNA-to-dsDNA ratio, and the RNA-to-dsDNA ratio. Each ratio was z-score standardized and tested as a predictor of recall in a separate linear mixed-effects model (lme4, v1.1-35.5; maximum likelihood)^111^ with sampling timepoint as an additional fixed effect and individual as a random intercept to account for repeated sampling. Fixed-effect p-values were obtained using Satterthwaite’s approximation (lmerTest, v3.1-3)^112^. P-values for the three indices were adjusted for multiple testing using the Benjamini-Hochberg procedure.

#### Holovirome profile generation

Per-sample holoviromes were constructed by combining paired VLP- and MGS-metaviromes into per-sample presence/absence profiles. The number of integrated prophages was estimated as the difference between the number of temperate phages detected in paired MGS- and VLP-metaviromes.

### Beta diversity and inter-sample similarity of the active virome

Bray-Curtis dissimilarities were computed on vOTU relative abundance profiles for all pairs of VLP-metaviromes using function vegdist from R package vegan (v2.6-8)^114^. PCoA was performed on Bray-Curtis dissimilarities using classical multidimensional scaling (cmdscale, stats R package v4.4.2^95^). The first two principal coordinates (PC1 and PC2) were visualized. For each timepoint, the average distance of infant samples to their group centroid in PCoA space was computed using vegan::betadisper^114^ on Bray-Curtis distances, as a measure of within-timepoint community heterogeneity.

Bray-Curtis similarities, calculated as 1 – Bray-Curtis dissimilarity, were compared between sample-pair categories (e.g., between infants versus between twins). The comparison was done using the same Wilcoxon rank-sum permutation procedure described above for the Jaccard analysis. Benjamini-Hochberg correction was applied across all contrasts.

The infant active virome was compared to maternal viromes using Bray-Curtis similarity. For each infant timepoint, two similarity values were retained: similarity to the infant’s own mother and mean similarity across all unrelated mothers. To assess whether similarity increased with infant age, a linear mixed-effects model was fitted per each mother type (own, unrelated), with infant age (months) as a fixed effect and individual infant as a random intercept: similarity ∼ infant age + (1|infant ID). To then test whether the age-related trajectory differed between mother types, a combined model with an age × mother type interaction was fitted (similarity ∼ infant age × mother type + (1|infant ID)). For visualization, per-timepoint means and their 95% bootstrap confidence intervals were estimated from 1,000 resamples.

### Virome composition and viral features association with phenotypes

Permutational multivariate analysis of variance (PERMANOVA; vegan::adonis2^114^, 999 permutations) was used to identify factors associated with the composition of the infant active virome (Bray-Curtis dissimilarities of vOTU relative abundances). The effect of subject identity was tested in a univariate model; infant age (timepoint) was tested in a separate model with infant identity included as a strata constraint to restrict permutations within individuals and account for repeated sampling. To assess the contribution of additional metadata to longitudinal virome composition listed in Supplementary Data 2, including feeding mode and delivery mode, separate PERMANOVAs were performed for each factor, with the phenotype and timepoint as terms, and infant identity included as a strata constraint. For cross-sectional phenotypes (constant within an infant; e.g., delivery mode), the analysis was restricted to infants with all four timepoints available, and permutations were constrained to exchange entire infants between phenotype groups while preserving the within-infant timepoint order. P-values were adjusted for multiple testing using the Benjamini–Hochberg procedure within hypothesis families separately for phenotypes hypothesized to shape virome composition and for health outcomes.

To identify the vOTUs driving the phenotype-associated differences in virome composition, vOTUs present in ≥ 10% of samples were tested for differential abundance using linear mixed-effects models (lme4::lmer^111^). For each vOTU, log-transformed abundance was modeled as a function of the phenotype and timepoint as fixed effects, adjusting for the percentage of reads lost during quality control and the percentage of aligned reads, sequencing batch, and non-bacterial sequence enrichment score (Virome QC^65^); infant identity was included as a random intercept. P-values were adjusted for multiple testing using the Benjamini–Hochberg procedure. Because only 2% and 13.3% of infant-detected vOTUs were classified at the species and genus levels, respectively, taxonomic-rank testing was restricted to the family rank. Viral families present in ≥10% of samples were tested using the same model as for vOTUs.

Viral features (e.g., viral richness and temperate phage abundance; full list in Supplementary Data 12) were tested against the same phenotypes using the same model framework. For normally distributed viral features (e.g., viral alpha diversity calculated as a Shannon index at the vOTU level), no log transformation was applied; a full list of applied models, pseudocounts, and transformations is provided in Supplementary Data 12. Feeding mode was tested for all viral features and timepoints, but visualized only for M1, M3, and M6 timepoints since very few infants (n = 12) were exclusively breastfed at M12. Viral features associated with virome-defining phenotypes (feeding or delivery mode) were adjusted for those phenotypes in the corresponding health outcome analyses. P-values were adjusted for multiple testing using the Benjamini–Hochberg procedure within hypothesis families separately for phenotypes hypothesized to shape virome composition and for health outcomes and for different types of viral features.

### Infant vOTU-sharing with mothers in the active virome

For each infant timepoint, the number of vOTUs shared with the infant’s own mother and with unrelated mothers was computed. For unrelated mothers, the mean number of shared vOTUs with all unrelated mothers was calculated. Sharing was then normalized by the number of vOTUs detected in the infant sample, yielding the proportion of the infant’s active virome shared with the maternal active virome. These proportions were modeled separately for related and unrelated mothers using a linear mixed-effects model (lme4::lmer^111^) with infant age as a fixed effect and individual infant as a random intercept.

To assess how phages predicted to infect different bacterial host genera contribute to overall infant–mother vOTU-sharing, a separate linear mixed-effects model (lme4::lmer) was calculated. The per-sample number of shared vOTUs assigned to the host genus was modeled as a function of sampling timepoint as a fixed effect and individual infant as a random intercept. Fixed-effect p-values were obtained using Satterthwaite’s approximation (lmerTest^112^).

### Determining the PPV in infants

PPVs were analyzed in infants with at least three longitudinal samples (n = 154). Cumulative richness over time was defined as the total number of unique vOTUs detected across all of an infant’s longitudinal VLP-metaviromes. The PPV was defined as the set of vOTUs present in at least three VLP-metaviromes of an infant, regardless of the age difference between timepoints (i.e., vOTUs detected in M3, M6, M12 or M1, M3, M6 were equally considered to be members of PPV). The TDV was defined as all vOTUs present in fewer than three VLP-metaviromes of an infant. Since both the cumulative richness over time and PPV size were lower in infants with three longitudinal VLP-metaviromes (n = 70) than in those with four (n = 84), these metrics were rarefied for the latter by subsampling to three timepoints and averaging across the four possible three-timepoint combinations. For PPV abundance estimation, no rarefaction was also applied and all PPV vOTUs identified per infant were used, as well as for the visualization of taxonomic and host distributions of PPV vOTUs pooled across infants.

For the expanded PPV definition, only vOTUs meeting all of the following criteria were considered: (1) temperate lifestyle; (2) detected at least once in a VLP-metavirome of the infant but assigned to its TDV; and (3) present in at least three of the infant’s holoviromes. The number of vOTUs meeting these criteria did not differ between infants with three and four timepoints and was therefore not rarefied.

To identify factors distinguishing PPV from TDV vOTUs (detection at M1, sharing with the mother, carriage of defense or anti-defense systems, and phage lifestyle), each factor was tested using a two-sided Fisher’s exact test. As each vOTU could appear as PPV in some infants and TDV in others, pair-level counts treated each (Infant, vOTU) combination as an independent unit of analysis. P-values across the four tests were adjusted for multiple testing using the Benjamini–Hochberg procedure.

The lifecycle dynamic of temperate phages was assessed in infants with all four timepoints available (n = 84) and restricted to temperate phages present in all four longitudinal holoviromes of an infant (n = 710). For each Infant × vOTU pair at each timepoint, the lifecycle state was classified as 1) “virion”, if the vOTU was detected in the infant’s VLP-metavirome, or 2) “prophage”, if the vOTU was detected only in the MGS-metavirome but not the VLP-metavirome. Based on the prophage/virion pattern across the four timepoints, each Infant × phage pair was assigned to one of five categories (all patterns shown in Figure 5e). These 16 longitudinal patterns were further binned into five categories: 1) stable prophage, if the vOTU was never detected in VLP-metaviromes; 2) induction, if the vOTU was detected as prophage at one or more initial timepoints and then as virion at all subsequent timepoints, with no reversion to prophage (e.g., prophage at M1, M3, M6, virion at M12); 3) stable replication, if the vOTU was detected in all four VLP-metaviromes; 4) lysogenization, if the vOTU was detected as virion at one or more initial timepoints and then as prophage at all subsequent timepoints, with no reversion to virion (e.g., virion at M1, M3, M6, prophage at M12); 5) switcher, if the vOTU did not fit any of the above categories.

### Data visualization

Results were visualized using a set of custom R scripts (R v4.4.2), including calls to functions from the packages ggplot2 (v4.0.0), dplyr (v1.1.4), tidyverse (v2.0.0), ggrepel (v.0.9.6), and patchwork (v.1.3.0). All boxplots were prepared using ggplot2 and represent standard Tukey type with IQR (box), median (bar) and Q1—1.5 × IQR/Q3 + 1.5 × IQR (whiskers).

### Statistics and reproducibility

All linear mixed-effects models were fitted with function lmer from lme4 R package, v1.1-35.5^111^. Fixed-effect p-values were obtained via Satterthwaite’s approximation as implemented in lmerTest v3.1-3^112^. To support reproducibility, complete statistical outputs, including model estimates, test statistics, and sample sizes, are provided in the corresponding Supplementary Data tables and test details are described in the corresponding Methods sections. For readability, Supplementary Data tables are cited once per paragraph for groups of related tests. Summary statistics tables were prepared using the R package skimr (v2.2.1).

### Data availability

All databases used for the dereplication of the NEXT virome catalog are publicly available and accessible by the links provided in Supplementary Data 31. The NEXT virome catalog is available at Figshare repository^115^ https://doi.org/10.6084/m9.figshare.32685267.

### Code availability

All scripts used in this study can be found at: https://github.com/GRONINGEN-MICROBIOME-CENTRE/Chiliadal_virome

## Acknowledgments

The data used in this manuscript were provided by LLNEXT. The authors are grateful to all participants of the LLNEXT cohort and thank all staff involved in participant recruitment, sample collection, and building and maintaining the cohort. We also thank Kate Mc Intyre for editing the manuscript and the UMCG Genomics Coordination Center and the Center for Information Technology of the University of Groningen for their support and for providing access to the Gearshift and Hábrók high-performance computing clusters. The authors further thank Hermie J.M. Harmsen and Erwin G.C. Raangs for providing the ML-II laboratory space for VLP extraction, Justin van Breen and Olga Minaeva for help with sample preparation for VLP extraction, and Marshellon Martinus for EGA data management.

The Lifelines NEXT cohort study received funds from the University Medical Center Groningen Hereditary Metabolic Diseases Fund, Health∼Holland (Top Sector Life Sciences and Health), the European Union, the Northern Netherlands Alliance (SNN), the provinces of Friesland and Groningen, the municipality of Groningen, Philips, and the Société des Produits Nestlé.

S.G. was supported by a scholarship from the Graduate School of Medical Sciences, University of Groningen, a grant from de Cock-Hadders Stichting (2021-08), and a Dutch Research Council (NWO) Rubicon grant (019.243EN.053). N.K. was supported by a grant from the de Cock-Hadders Stitchting (2026-81). A.F-P. was supported by a grant from the de Cock-Hadders Stichting (2023-50). T.S. was supported by a grant from EASI-Genomics and Corundum Systems Biology. E.R.W. received funding from a UK Research and Innovation grant under the UK Government’s Horizon Europe funding guarantee EP/X030377/1, the Biotechnology and Biological Sciences Research Council BB/X003051/1, the Biotechnology and Biological Sciences Research Council/National Science Foundation research grant 2321502, and the 2020 Philip Leverhulme Prize in Biological Sciences. J.F. is supported by a European Research Council (ERC) Consolidator grant (grant agreement No. 101001678), an NWO VICI grant (VI.C.202.022), an NWO KIC grant (KICH1.LWV04.21.013), the AMMODO Science Award 2023 for Biomedical Sciences from Stichting Ammodo, and the Dutch Heart Foundation AtheroNeth project (01-001-2024-0601). A.K. is supported by UMCG Startersbeurs grant O/695834. A.K. and A.Z. are supported by the NWO Gravitation grant ExposomeNL 024.004.017. A.Z. is supported by NWO-VICI grant VI.C.232.074, NWO KIC grant KICH1.LWV04.21.01, ERC Starting Grant 715772, ZonMW ME/CFS grant 10091012110015, EU Horizon Europe Program grant INITIALISE (101094099), and EU Horizon Europe Program grant DarkMatter (“ID-DarkMatter-NCD” (project number 101136582)).

## Author information

Lifelines NEXT cohort study

Milla F. Brandão-Gois^1^, Marcel Bruinenberg^2^, Siobhan Brushett^1,3^, Jackie A.M. Dekens^1,4^, Sanzhima Garmaeva^1^, Sanne J. Gordijn^5^, Soesma A. Jankipersadsing^1^, Ank de Jonge^6,7^, Trynke R. de Jong^2^, Gerard H. Koppelman^8,9^, Marlou L.A. de Kroon^3^, Folkert Kuipers^8,10^, Alexander M. Kurilshikov^1^, Lilian L. Peters^6,7^, Jelmer R. Prins^5^, Sijmen A. Reijneveld^3^, Sicco Scherjon^5^, Jan Sikkema^4^, Trishla Sinha^1^, Johanne E. Spreckels^1^, Aline B. Sprikkelman^8,9^, Morris A. Swertz^1^, Henkjan J. Verkade^8^, Cisca Wijmenga^1^, Alexandra Zhernakova^1^

^1^Department of Genetics, University of Groningen, University Medical Center Groningen, Groningen, the Netherlands

^2^Lifelines Cohort Study and Biobank, Groningen, the Netherlands

^3^Department of Health Sciences, University of Groningen, University Medical Center Groningen, Groningen, the Netherlands

^4^Innovation Center, University Medical Center Groningen, Groningen, the Netherlands

^5^Department of Obstetrics and Gynecology, University of Groningen, University Medical Center Groningen, Groningen, the Netherlands

^6^Department of Primary and Long-term Care, University of Groningen, University Medical Center Groningen, Groningen, the Netherlands

^7^Midwifery Science, Amsterdam University Medical Center, Vrije Universiteit Amsterdam, AVAG, Amsterdam Public Health, Amsterdam, the Netherlands

^8^Department of Pediatrics, University of Groningen, University Medical Center Groningen, Groningen, the Netherlands

^9^Groningen Research Institute for Asthma and COPD (GRIAC), University of Groningen, University Medical Center Groningen, Groningen, the Netherlands

^10^European Research Institute for the Biology of Ageing (ERIBA), University of Groningen, University Medical Center Groningen, Groningen, the Netherlands

## Contributions

S.G. and A.Z. conceptualized and supervised the study. S.G., T.S., J.E.S., S.B., L.L.N., A.K., and A.Z. initiated, designed and supported the LLNEXT cohort study. A.Z. acquired funding. S.G., T.S., J.E.S., S.B., and L.L.N gathered and processed the biological data. S.G., A.F.P., T.S., J.E.S., S.B., and L.L.N processed the phenotypic data. S.G., A.F.P., T.S., J.E.S., S.B., and A.K. processed the metagenomic data. S.G., N.K., S.S., J.G-A., and M.K. extracted and generated VLP-metaviromes data. S.G., A.F.P., and A.G. developed the assembly-based viral discovery pipeline. N.K. developed auxiliary genes annotation pipeline. S.G., N.K., and A.F.P. performed the data analysis. S.G. and N.K. performed the statistical analysis. A.G., J.A.D.D, E.R.W., J.F., and A.K. supported and discussed the data analysis. S.G., N.K., and A.Z. wrote the manuscript and created the figures. All authors have read and agreed to the published version of the manuscript.

## Competing interests

A.Z. received a speaker fee from Nestlé and AVOLA. Other authors declare no competing interests. The funders had no role in study design, data analysis, data interpretation, writing of the manuscript, and the decision to publish.

---

**Supplementary Figure 1.**
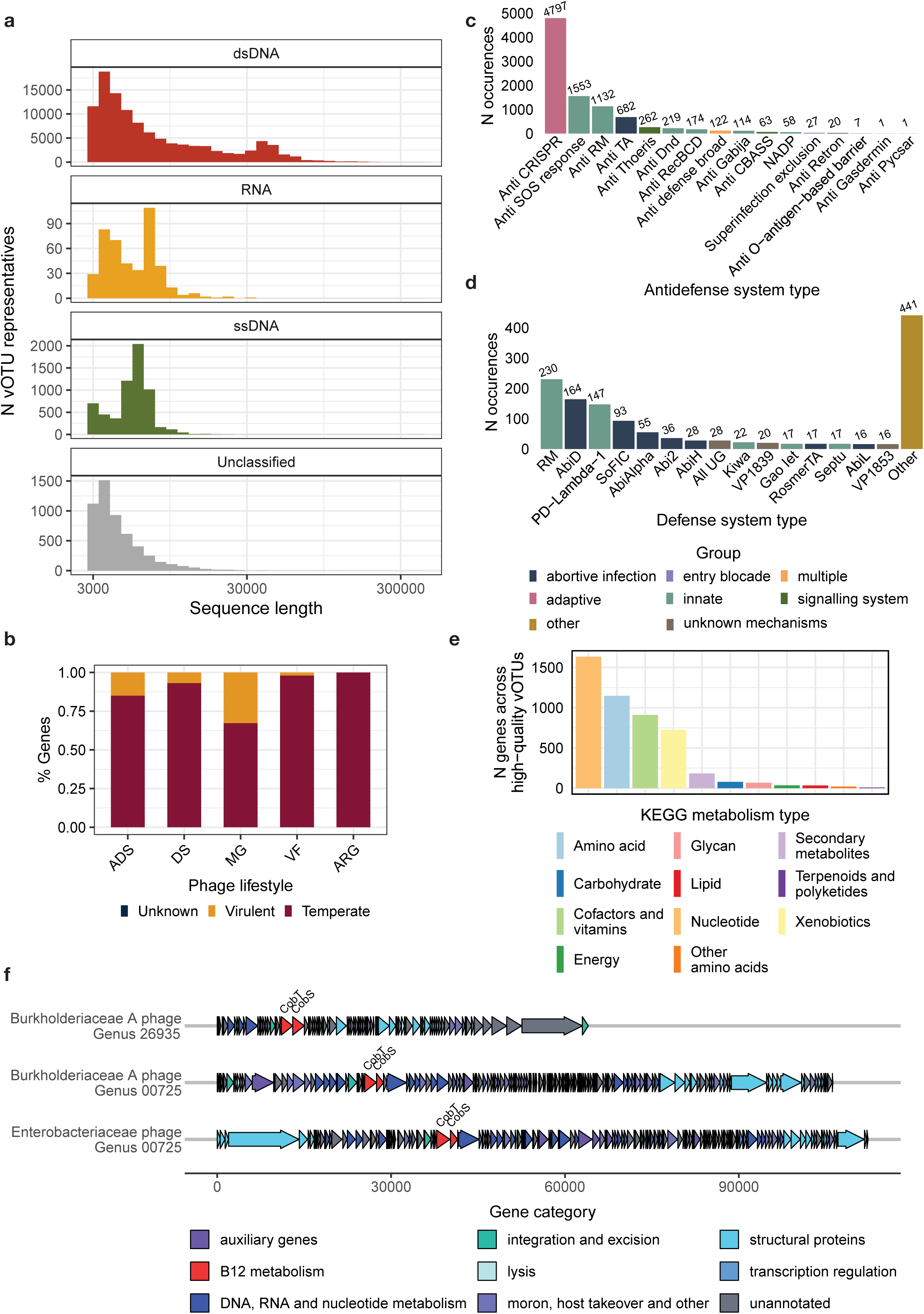
Length distribution of vOTU representatives and characteristics of auxiliary genes encoded by the viruses. a. Genome length distribution of vOTUs in the NEXT virome catalog, faceted by genome type (dsDNA, ssDNA, RNA, Unclassified). X-axis shows genome length (bp). Y-axis shows the number of vOTUs. b. Distribution of auxiliary genes in phages stratified by predicted lifestyle. Stacked bars show the relative proportion of systems (ADS, DS) or genes (MGs, VFs, ARGs) within each auxiliary category. ADS, anti-defense systems; MG, metabolic genes; DS, defense systems; VF, virulence factors; ARG, antibiotic resistance genes. c, d, Distribution of (c) anti-defense systems and (d) defense systems across HQ vOTUrs in the NEXT virome catalog. Bar heights represent the total number of detections per system, with exact counts indicated above each bar. Bars are colored according to manually curated groupings based on the prokaryotic defense mechanism (c, targeted by the vOTU; d, carried by the vOTU), following a previously described classification framework^55^. e. Distribution of MGs identified in HQ vOTUrs. Bar heights indicate the total number of genes assigned to each KEGG metabolism group, with bars color-coded according to KEGG metabolism groups. f. Gene maps of the selected HQ vOTUrs encoding the CobS and CobT genes involved in vitamin B12 metabolism. Genes are color-coded according to functional categories as follows: genes annotated as “tail”, “connector”, or “head and packaging” by PHROG are grouped into the “structural proteins” category; genes annotated as “moron, auxiliary metabolic gene and host takeover” or “other” by PHROG are grouped into the “moron, host takeover, and other” category; genes identified as auxiliary genes using the targeted pipeline are grouped into the “auxiliary genes” category; other genes annotated by PHROG are assigned to the original PHROG-coded category. CobS and CobT genes are marked within each genome.

**Supplementary Figure 2.**
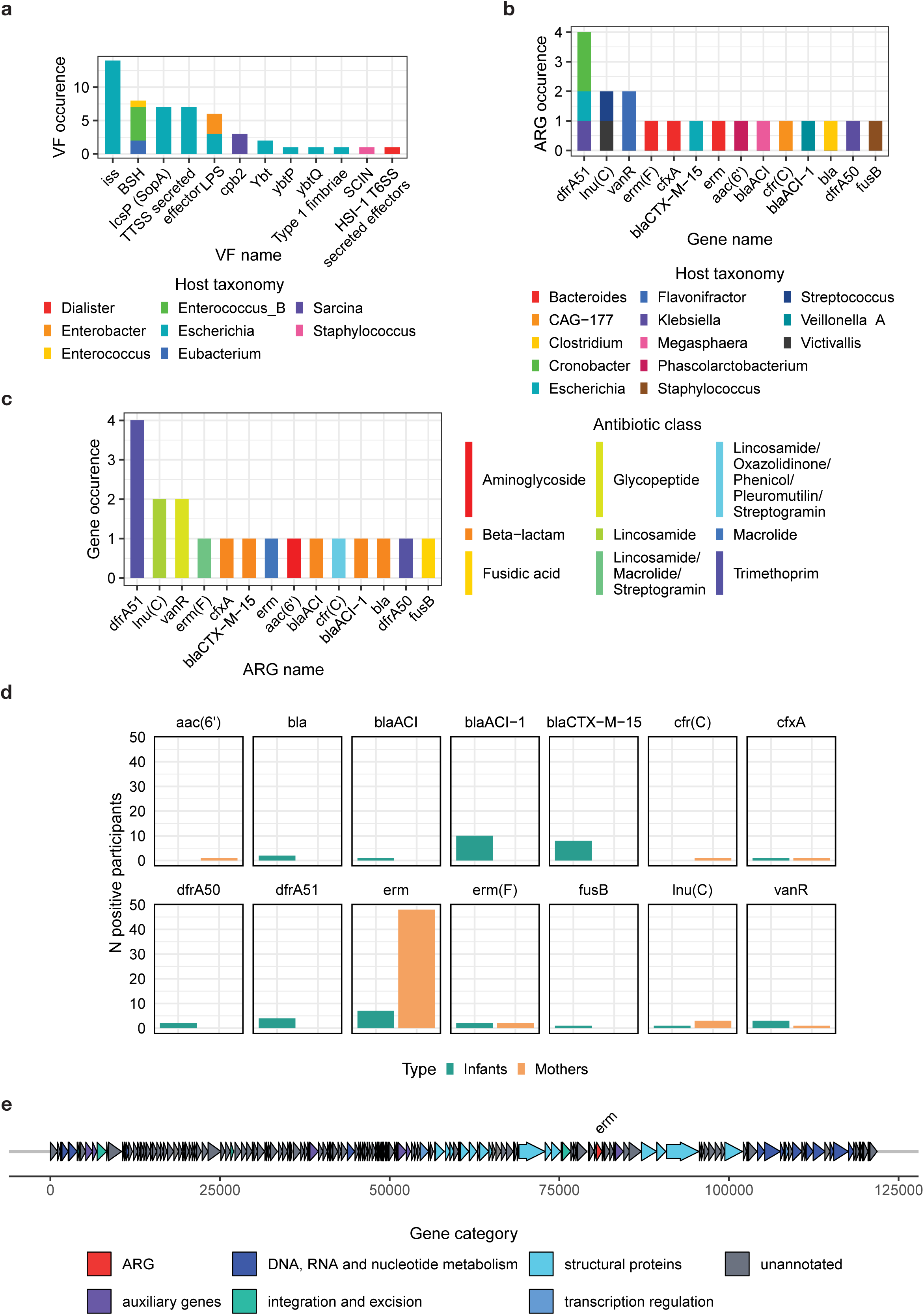
Virus-encoded VF and ARG distributions and characteristics. a-c. Occurrence of (a) VF genes and (b, c) ARG across HQ vOTUrs. Y-axis shows the number of detections across vOTUrs. X-axis shows curated gene names from (a) the VFDB or AMRFinderPlus databases or (b, c) AMRFinderPlus only. Bars are colored according to (a, b) the predicted bacterial host genus of the virus carrying the gene or (c) the antibiotic class defined in AMRFinderPlus. d. Prevalence of vOTUrs encoding ARGs, stratified by sample ARG, and color-coded as infants (green) and mothers (orange). ARG names are shown above each facet and correspond to entries in the AMRFinderPlus database. Bar heights indicate the number of individuals positive for at least one virus encoding the corresponding ARG e. Gene map of a Bacteroides phage encoding a 23S rRNA methyltransferase Erm (erm) gene. Gene annotations were assigned using the PHROG database, and ARG was identified with AMRFinderPlus. Genes are color-coded according to the following functional categories: “structural proteins” = genes annotated as “tail”, “connector”, or “head and packaging” by PHROG and “auxiliary genes” = genes identified as auxiliary genes using the targeted pipeline. Other genes annotated by PHROG are assigned to the original PHROG-coded category. 23S rRNA methyltransferase Erm is assigned to the ARG category.

**Supplementary Figure 3.**
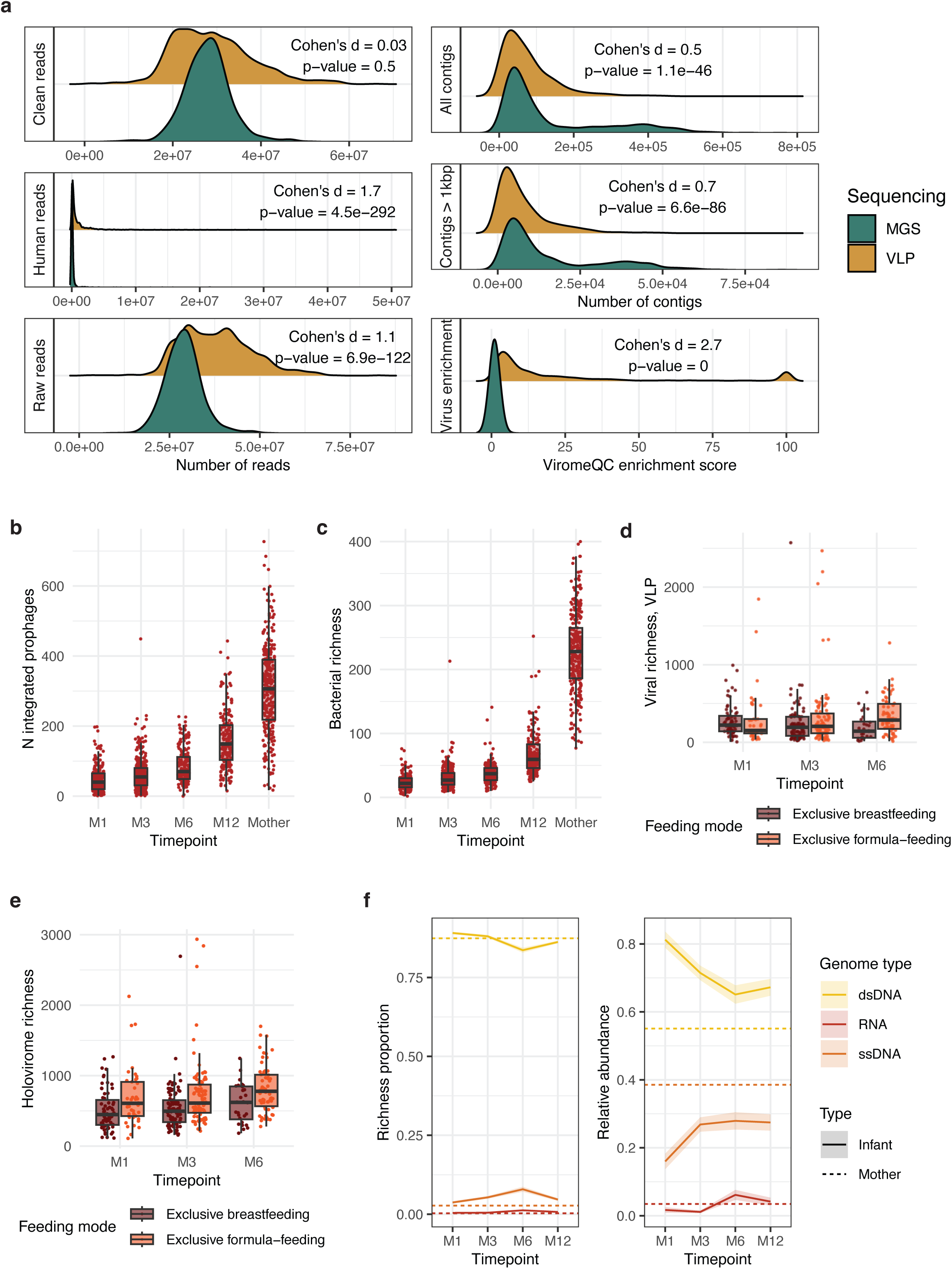
Metavirome characteristics and features of virome by host, lifestyle, and early-life exposures. a. Ridgeline plots of genomic characteristics of MGS- and VLP-metaviromes. Each ridge shows the distribution of one characteristic across samples within each metavirome type. Statistical comparisons between MGS and VLP were performed using a LMM. b. Number of integrated prophages per sample in infants across timepoints and mothers. Integrated prophages were defined as the difference between the number of temperate phages detected in the holovirome and the number detected in the VLP-metavirome (Methods). c. Number of bacterial species (species-level genome bins (SGB) level) identified in MGS samples in infants across timepoints and mothers. d, e. Viral richness in VLP (d) and holovirome richness (e) in infants across timepoints (M1, M3, M6) by feeding mode. f. Richness and summed relative abundance of vOTUs by virus genome type in the infant active virome across timepoints. The mean maternal level is indicated as a single dashed horizontal line per facet. All boxplots show the median (center line), 25^th^ and 75^th^ percentiles (hinges), and whiskers extending to 1.5× the interquartile range.

**Supplementary Figure 4.**
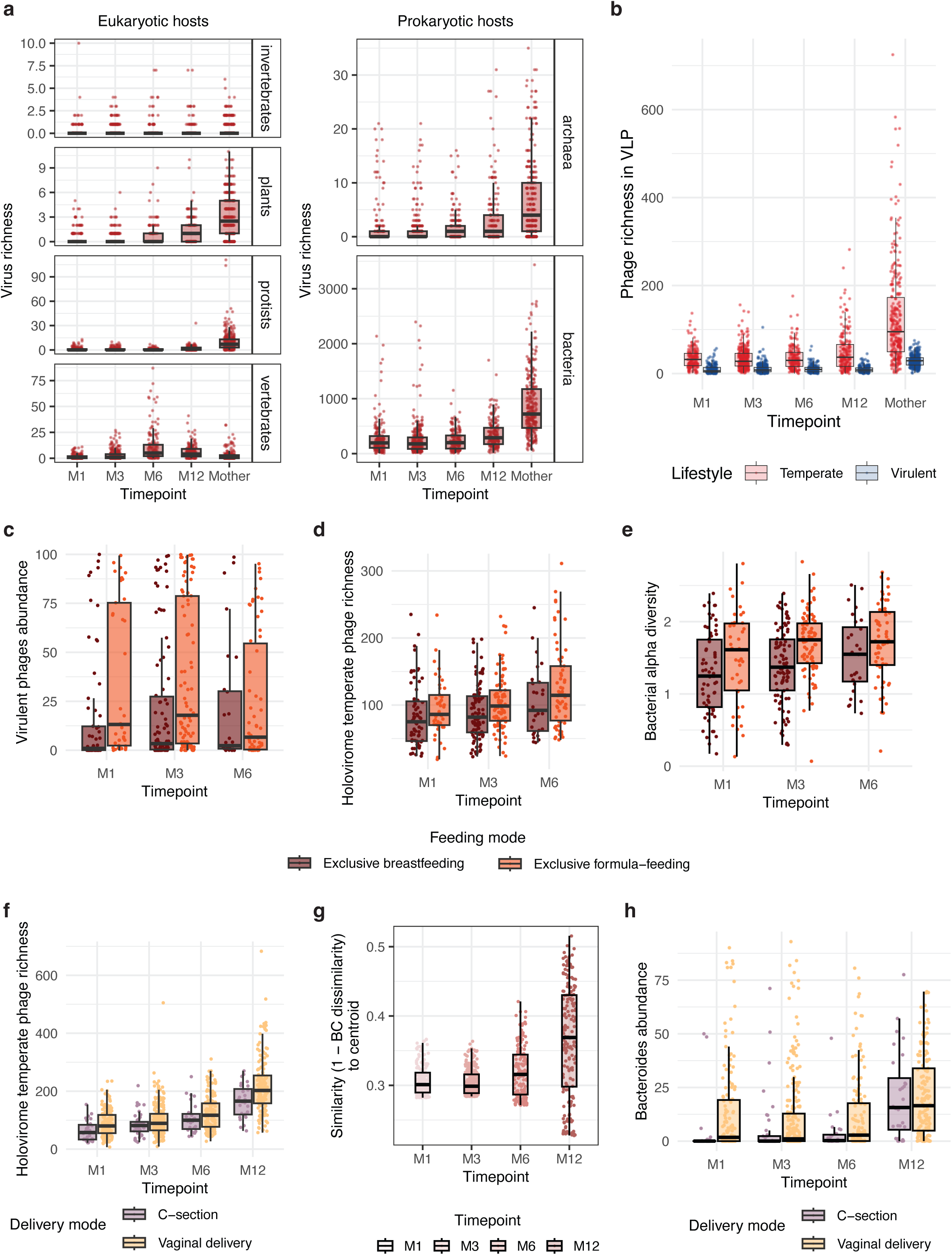
Temporal dispersion of the infant active virome and its modulation by early-life exposures and health outcomes. a. Temporal dynamics of vOTU richness in the infant and maternal active virome by predicted host, faceted by general host category (eukaryotic vs prokaryotic). b. Phage richness in the active virome across infant timepoints and mothers, colored by predicted phage lifestyle. c-e. Virulent phage abundance (c), temperate phage richness in holovirome (d), and bacterial alpha diversity at SGB level (e) in infants across timepoints (M1, M3, M6) by feeding mode. f. Temperate phage richness in the holovirome across infant timepoints by delivery mode. g. Pe r-sample similarity (1 − Bray-Curtis dissimilarity) to the timepoint centroid across infant timepoints, visualizing within-group dispersion. Points and boxplots are colored by timepoint along a temporal gradient. h. Relative abundance of the *Bacteroides* genus in infants across timepoints, colored by delivery mode. All boxplots show the median (center line), 25^th^ and 75^th^ percentiles (hinges), and whiskers extending to 1.5× the interquartile range.

**Supplementary Figure 5.**
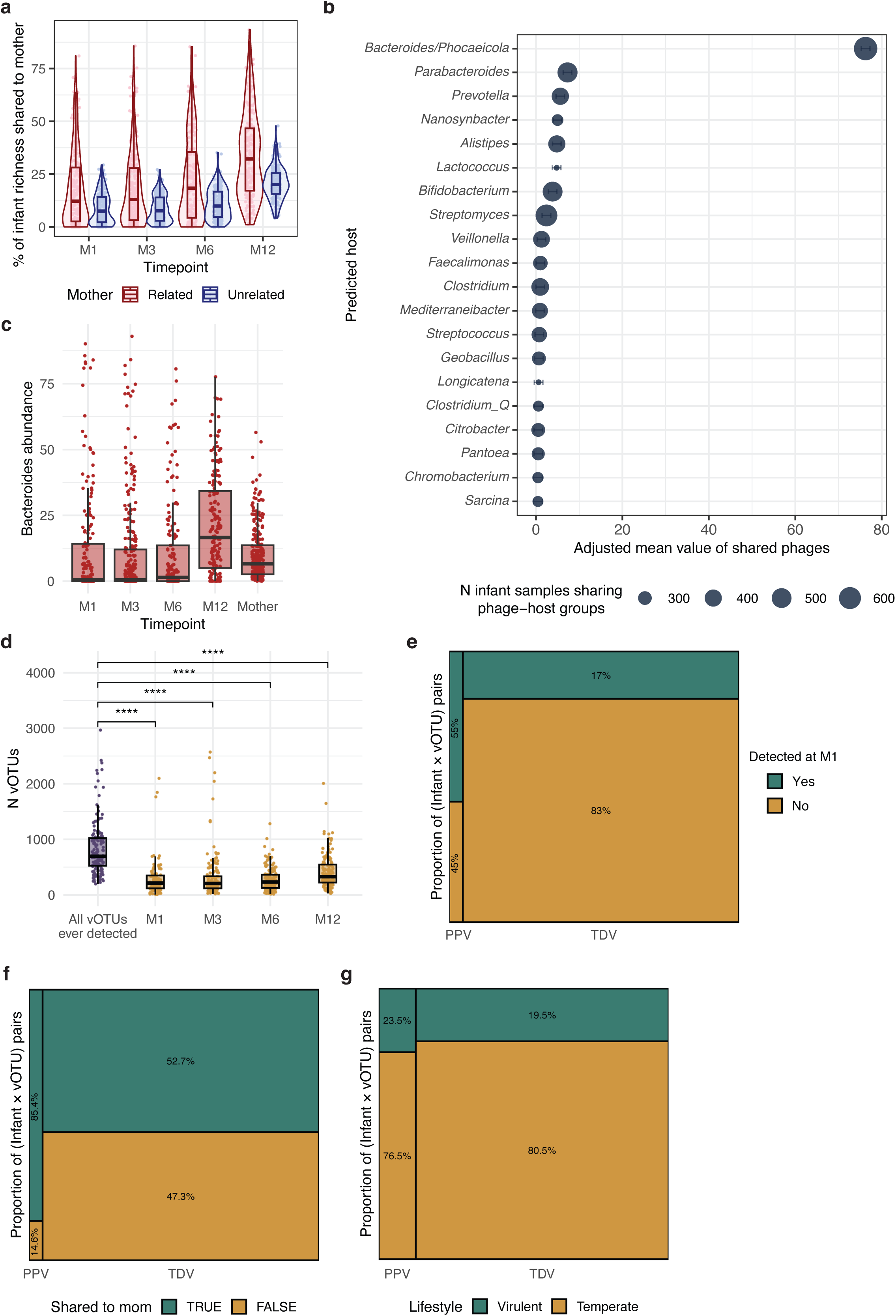
Persistence and mother–infant sharing of the infant active virome. a. Percentage of infant vOTU richness in the active virome shared with mothers across infant timepoints, colored by mother kinship. Each dot represents an infant timepoint sample paired with its own (related) or unrelated mother (mean). b. Per-sample contribution of predicted bacterial hosts to vOTUs shared between infants and mothers in the active virome. Each point represents the timepoint-adjusted estimated mean contribution for a host genus, with horizontal bars showing 95% confidence intervals. Point size scales with phage-host groups prevalence across infant samples. Only host genera with top-20 prevalent phage-host groups are shown. c. Relative abundance of the *Bacteroides* genus in infants across timepoints and mothers. d. Number of vOTUs detected per sample in the infant active virome across timepoints, alongside the cumulative number of vOTUs detected across all timepoints per infant (rightmost boxplot). Each dot in the per-timepoint boxplots represents one infant timepoint sample. Each dot in the cumulative boxplot represents one infant. Each timepoint was compared to the cumulative richness using Tukey’s post-hoc test. **** FDR < 0.05. All boxplots show the median (center line), 25^th^ and 75^th^ percentiles (hinges), and whiskers extending to 1.5× the interquartile range. e-g. Mosaic plots of infant vOTUs in the active virome, pooled across infants and segmented along two dimensions: PPV versus TDV status and (e) detection at M1, (f) sharing with own mother, (g) vOTU lifestyle Tile area is proportional to the number of vOTUs in each combination.

## Notes

https://doi.org/10.6084/m9.figshare.32685267

https://github.com/GRONINGEN-MICROBIOME-CENTRE/Chiliadal_virome

### Competing Interest Statement

A.Z. received a speaker fee from Nestle and AVOLA. Other authors declare no competing interests. The funders had no role in study design, data analysis, data interpretation, writing of the manuscript, and the decision to publish.

